# Integrating gene expression, mutation and copy number data to identify driver genes of recurrent chromosome-arm losses

**DOI:** 10.1101/2025.02.22.639659

**Authors:** Ron Saad, Ron Shamir, Uri Ben-David

## Abstract

Aneuploidy is a hallmark of cancer, yet the specific genes driving recurrent chromosome-arm losses remain largely unknown. Here, we present a systematic framework integrating gene expression, mutation, and copy number data to identify candidate driver genes of cancer type-specific recurrent chromosome-arm losses across 20 cancer types, using ∼7,500 tumors from The Cancer Genome Atlas. By analyzing focal deletions and point mutations that co-occur with, or are mutually exclusive with, chromosome-arm losses, we pinpoint 311 candidate drivers associated with 160 cancer type-specific recurrent events. Our approach identifies known aneuploidy drivers such as *TP53* and *PTEN*, while revealing multiple novel candidates, including established tumor suppressor genes not previously linked to aneuploidy. Furthermore, we leverage gene expression changes associated with these chromosome-arm losses to propose pathway-level alterations that may drive cancer progression. Integrating these findings highlights key candidate drivers underlying the observed gene expression alterations, thereby reinforcing their biological relevance. This work provides the first comprehensive catalogue of candidate driver genes for recurrently lost chromosome-arms in human cancer.

## Introduction

Aneuploidy, an abnormal number of chromosomes or chromosome arms (hereafter referred to as ‘arms’) in a cell, has long been recognized as a hallmark of cancer [1]. Despite being very common in cancer, occurring in ∼90% of solid tumors [2], its role in cancer initiation and progression is still not fully understood [3].

Advances in molecular biology research tools, such as high-throughput sequencing and CRISPR, along with improved data collection efforts through comprehensive databases like The Cancer Genome Atlas (TCGA) [4] and Cancer Cell Line Encyclopedia [5], have significantly enhanced our ability to characterize aneuploidy in cancer. These developments have provided deeper insights into the complex landscapes of chromosomal alterations in cancer [2], and into the selection pressures that shape them [1], [3], [6], [7], [8], [9], [10], [11], [12].

An important observation made in recent years is that the role of specific alterations is highly context-dependent, influenced by factors such as tumor stage, cell type, and interactions with the immune system [1], [3]. These factors contribute to the selection pressures that shape the complex landscape of aneuploidy in cancer. We and others have recently demonstrated the importance of negative selection in shaping the aneuploidy landscapes of human cancer [6], [13]. Nonetheless, the recurrence of particular alterations within a given context suggests that these recurrent changes are positively selected, indicating their potential role in driving cancer initiation and progression [3]. Importantly, however, the genes that drive common aneuploidies (hereafter referred to as ‘drivers’) remain poorly characterized, with only a handful of cases in which candidate driver genes have been demonstrated to underlie the recurrence of a specific aneuploidy [10], [11], [14]. Even when specific strong drivers were identified, whether additional genes on the same arm also contribute to aneuploidy recurrence remains largely unknown [3].

Identifying the elements that underlie the positive selection for specific recurring aneuploidies is highly challenging for several reasons [3]. First, each such event impacts hundreds or thousands of genetic elements, making it difficult to pinpoint the actual drivers. Every arm harbors multiple tumor-suppressor genes (TSGs) and oncogenes, but not every cancer-related gene would necessarily be the driver of an arm-level copy number alteration. Second, the effects of these alterations are highly context-dependent, making them difficult to study and to model accurately. Third, generating cancer models with specific aneuploidies remains a significant technical challenge, limiting our ability to investigate the effects of these events comprehensively.

Previous studies in this field have focused on identifying characteristics of arms that influence their likelihood of being deleted or amplified. For instance, Davoli et al. [15] demonstrated that the density and potency of TSGs are positively correlated with the frequency of arm loss, and negatively correlated with the frequency of its gain, while oncogene density and potency exhibited the opposite trends. We recently corroborated these findings and further identified compensation by paralogs as being positively correlated with an arm’s loss frequency [6]. While these studies have provided valuable insights into the relationship between arm features and aneuploidy prevalence, they do not directly address the identification of specific drivers of these events. A recently developed tool, BISCUT [13], begins to address this challenge by analyzing telomere- and centromere-bound copy-number event distributions. Notably, the systematic approach described in[13] was strictly based on copy-number analyses. We therefore speculated that integrating other genomic modalities, namely point mutations and gene expression, could further improve the identification of candidate aneuploidy driver genes.

In this work, we propose a new framework for identifying driver genes of cancer type-specific recurring arm losses. Our analysis leverages mutation, gene expression, and copy number data from ∼7,500 tumor samples across 20 cancer types in the TCGA dataset. We examine the relationship between arm loss events and other genomic alterations that can inactivate a given gene, specifically point mutations and focal deletions. These analyses allowed us to identify genes frequently affected both by arm losses and by other mechanisms of gene inactivation, suggesting that they function as tumor suppressors whose bi-allelic inactivation involves an arm loss. We also identified genes predominantly affected by a single mechanism, either an arm loss or a focal event but not both. This may indicate that their mono-allelic inactivation provides a selective advantage to cancer, though their bi-allelic loss may not, or that the combined effect of the arm loss and the focal event on closely residing genes is detrimental. Lastly, we used gene expression data to identify changes in pathway activity associated with recurrent arm losses and link these alterations to our candidate driver genes in order to further refine our list of arm loss driver genes.

We identified known drivers of specific common aneuploidies, such as *TP53* [16], [17], [18] and *PTEN* [19], [20], [21], and also nominated new candidates, overall proposing 311 drivers for 160 cancer type-specific recurring arm losses. Some of these novel candidates are established TSGs not previously associated with aneuploidy, whereas others do not have an established role in human tumorigenesis. We therefore provide the first comprehensive resource of candidate driver genes for all recurrently lost arms in human cancer. Experimental validations will be required to demonstrate the driving role of the newly identified candidate genes.

## Results

### Approach overview

A schematic representation of our approach for identifying cancer type-specific driver genes associated with recurring arm losses is shown in **Fig. 1a**. We first calculated the prevalence of loss of each arm across 20 cancer types, using ∼7,500 samples from TCGA. We defined an arm loss as **common** in a specific cancer type if it was lost in >20% of the samples. This analysis identified 230 common arm losses across cancer types, hereinafter referred to as chromosome arm-cancer type (CA-CT) pairs (top left panel). For each arm and cancer type in the identified CA-CT pairs, we conducted expression and genetic analyses to identify potential driver genes and uncover the consequences of arm loss. The expression analysis was done by comparing samples with the arm loss to those without it (bottom left panel). The genetic analysis was done by using mutations and copy number data (bottom right panel). In order to identify drivers of arm loss, we defined four **perturbation patterns**: (1) Genes frequently affected by a focal deletion when the arm is lost (FD+AL). (2) Genes frequently affected by a point mutation when the arm is lost (PM+AL). (3) Genes frequently affected by focal deletion only when the arm is not lost (FD-AL). (4) Genes commonly affected by a point mutation only when the arm is not lost (PM-AL). Finally, we summarized the expression and genetic results for each arm across cancer types and perturbation patterns (top right panel).

**Figure 1:**
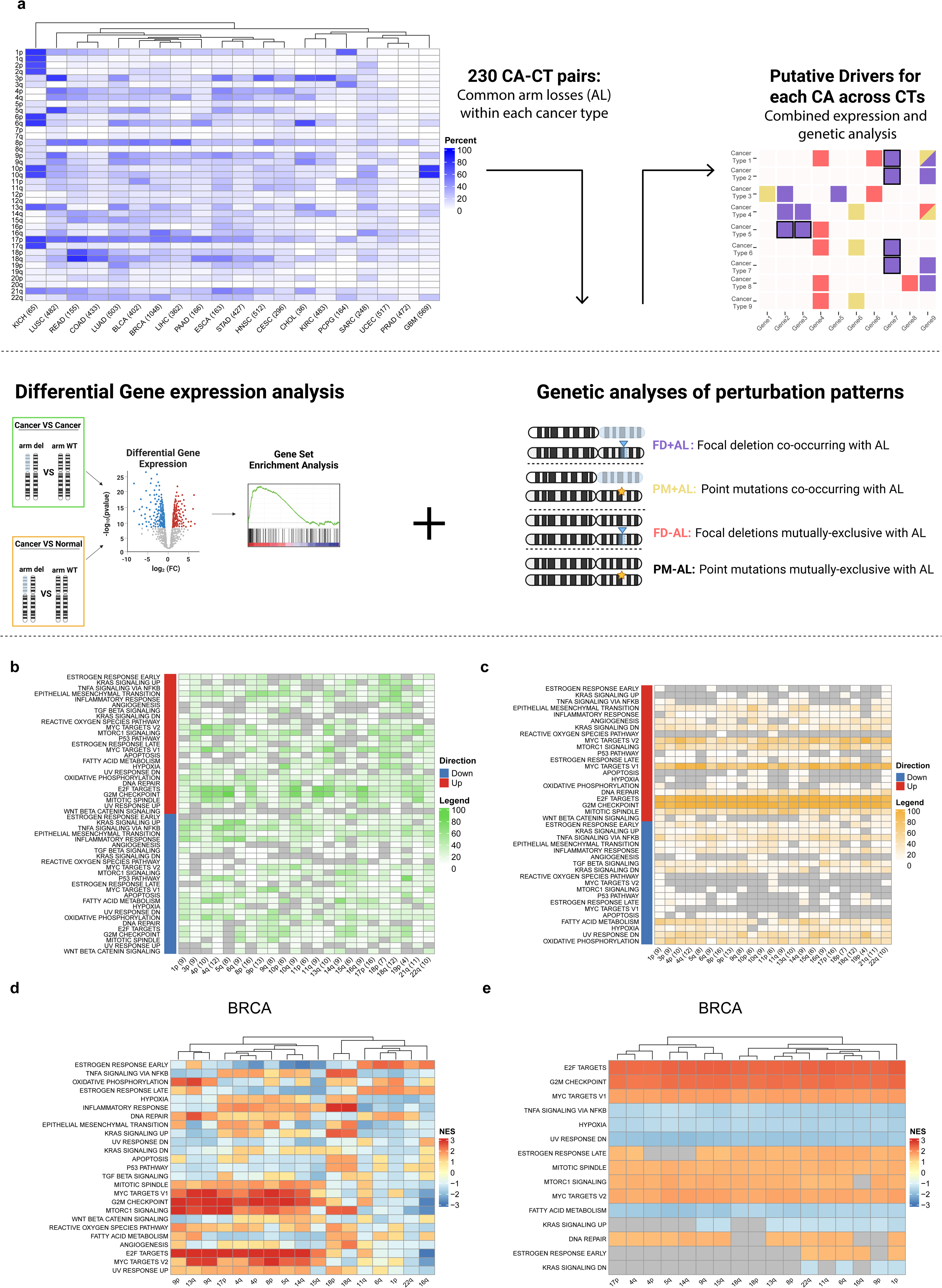
Approach overview and gene expression analysis. **a. Overview of the analysis methodology.** We calculated the prevalence of loss of each arm across 20 cancer types in ∼7,500 TCGA samples and identified 230 arms that were commonly lost in specific cancer types (top left). The expression changes associated with each such arm loss were studied in the corresponding cancer type (bottom left), to identify pathways associated with the arm loss. Genetic analysis was performed on each such arm in the corresponding cancer type (bottom right), to nominate drivers of the arm loss. The results of these analyses across cancer types were then integrated and visualized (top right). **b. + c. Pathway dysregulation associated with recurrent chromosome-arm losses across cancer types.** For each arm the percentage of cancer types in which a given Hallmark gene set is significantly up or down regulated out of all cancer types showing a common loss of the arm was calculated, comparing tumors with vs. without each arm loss (b) or tumors with the arm loss vs. normal tissues (c). 25 cancer-related gene sets are visualized. Numbers in parentheses are the number of cancer types in which the arm is commonly lost. **d. + e. Pathway dysregulation associated with recurrent chromosome-arm losses in BRCA**. For each BRCA recurrent arm loss, the upregulation or downregulation of cancer-related Hallmark gene sets are shown, comparing tumors with vs. without each arm loss (d) or tumors with the arm loss vs. normal tissues (e). NES: GSEA normalized enrichment score. BRCA, breast cancer.

### Gene expression changes associated with recurrent arm losses

To identify the gene expression consequences of specific arm losses in each cancer type, we performed the following comparisons for each CA-CT pair (**Fig. 1a**, bottom left panel): (1) Within-cancer: cancer samples that harbor the arm loss were compared to cancer samples without the loss. (2) Between cancer and normal tissue: cancer samples that harbor the arm loss were compared to the normal-adjacent tissue samples.

For each comparison, we performed a differential gene expression (DGE) analysis followed by gene set enrichment analysis (GSEA) [22] using the MSigDB ‘Hallmark’ gene sets collection[23] (**Supp. Table 1**). This collection includes 50 gene sets that represent well-defined biological states or processes, offering a high-level perspective on the differences between groups in each comparison. Our preliminary within-cancer type comparisons showed consistent patterns across different arms. We hypothesized that these findings reflect the general effects of high aneuploidy levels. To account for this, we repeated the analysis controlling for the confounder effect of aneuploidy levels in these comparisons using IPTW (see Methods). On average, 19 and 17 Hallmark gene sets were differentially expressed between the aneuploid tumors and the non-aneuploid tumors or the normal adjacent tissues, respectively (**Supp. Fig. 1**). It is important to note that the sample sizes of normal tissues were considerably smaller than those of the tumor groups, limiting the statistical power for comparisons involving normal samples. While these results suggest that arm losses may indeed have strong transcriptional consequences, the observed gene expression changes were likely influenced by additional factors associated with the recurrent arm losses, complicating the identification of the true gene expression consequences of these aneuploidies.

Next, we sought to determine whether a specific arm loss is associated with consistent gene expression changes across cancer types. For each recurrent arm loss, we calculated the fraction of cancer types in which each gene set was significantly up- or down-regulated (**Fig. 1b,c**). When analyzing the results of the within-cancer type comparisons, we found that, in most cases, the gene-expression changes associated with an arm loss were not ubiquitous across cancer types (**Fig. 1b** and **Supp. Table 1**). In contrast, when analyzing the cancer vs. normal tissue comparisons, the effects were consistent across arms (**Fig. 1c** and **Supp. Table 1**), indicating that these comparisons were likely dominated by general effects of cancer.

Finally, we evaluated the similarity of the gene expression changes that are associated with different arm losses within each cancer type (shown for BRCA in **Fig. 1d,e**; and for all other tumor types in **Supp. Fig. 2** and **Supp. Fig. 3**). In the within-cancer type comparisons, many of the recurrent arm losses were associated with the same gene expression changes (**Fig. 1d**). The clustering of the arms into a few groups with similar effects likely reflects the co-occurrence of arm losses [7], as well as general chromosome-independent effects of aneuploidy (or other aneuploidy-associated genomic alterations) [24], [25], [26]. The cancer vs. normal tissue comparisons showed that the same pathways were associated with nearly all recurrent arm losses, further indicating that the observed effects were largely influenced by general cancer-related gene expression changes (**Fig. 1e**).

Given the limitations of the gene expression analyses, we concluded that associating recurrent arm losses with dysregulated gene expression pathways, albeit informative, is not sufficient in and of itself for identifying and prioritizing potential drivers of these aneuploidies.

### Focal deletions co-occurring with arm losses

We next turned to a genetic analysis. We began by identifying focal deletions on a particular arm that frequently co-occur with the loss of that arm (pattern FD+AL, **Fig. 2a**). For each CA-CT pair, we applied GISTIC2.0 [27], a tool designed to identify regions significantly amplified or deleted across a set of samples, to samples harboring the arm loss, identifying 118 focal deletions across 99 pairs. These deletions result in bi-allelic inactivation of specific chromosomal regions. We found that most genes were rarely affected by both focal and arm-level losses simultaneously (shown for BRCA in **Fig. 2b** and for all other tumor types in **Supp. Fig. 4**). The regions that are frequently co-deleted by both an arm loss and a focal deletion are rather small (median size of ∼350Kb, encompassing a median of 4 genes), consistent with a strong negative selection against the bi-allelic loss of large chromosomal regions [28]. In regions with three or fewer genes, we nominated the genes with the highest deletion prevalence as candidate drivers of the combined focal and arm losses. For larger regions, we nominated candidate drivers only if they appeared in a small region in other cancer types (see Methods). It is important to note that we only considered protein-coding genes, although the recurring loss of the region may be driven by a non-coding genetic element.

**Figure 2:**
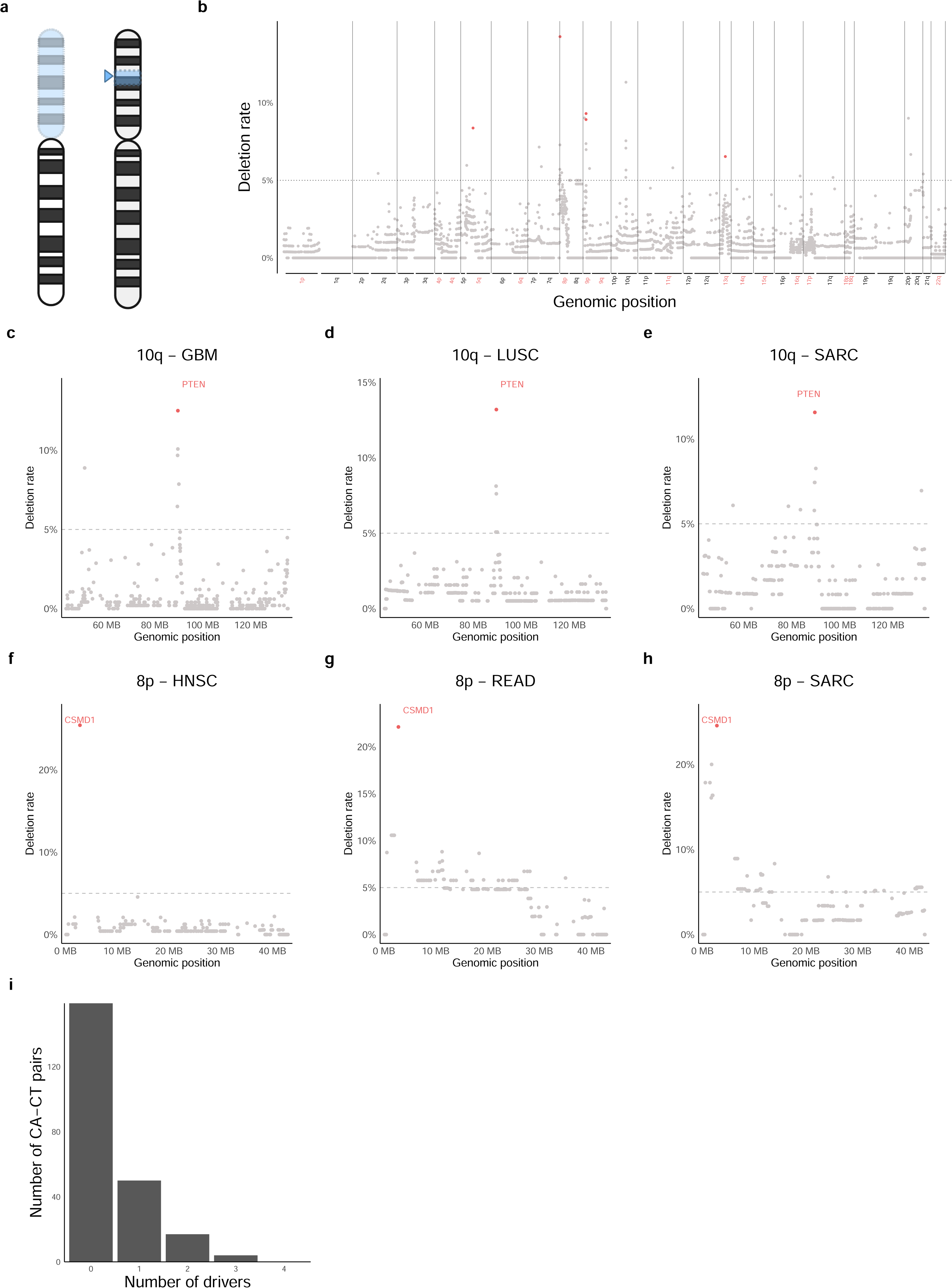
Focal deletions co-occurring with arm losses. **a. Schematic representation of the FD+AL inactivation pattern.** **b. The prevalence of focal deletions co-occurring with arm losses in BRCA**. The focal deletion rate of each gene when the other copy of the arm on which it resides is lost in BRCA samples. Arms commonly deleted are colored in red. Genes identified as drivers by this pattern are colored in red. **c. + d. + e. The prevalence of focal deletions co-occurring with Chr10q loss in GBM, LUSC and SARC**. **f + g. + h. The prevalence of focal deletions cooccurring with Chr8p loss in HNSC, READ and SARC**. **i. Distribution of the number of drivers identified in CA-CT pairs for the FD+AL pattern.**

The full list of genes that were identified in this pattern analysis is provided in **Supp. Table 2**. A prominent example is observed in Chr10q in multiple cancer types, including GBM, LUSC, and SARC (**Fig. 2c-e** and **Supp. Fig. 5**). In this case, the bi-allelic deletions affect small regions containing only a few genes, and our analysis identified the known TSG *PTEN* as the culprit of this arm loss. Indeed, *PTEN* was previously reported to drive the loss of Chr10q in GBM and other tumor types [19], [20], [21].

Another interesting example is observed in arm Chr8p (**Fig. 2f-h** and **Supp. Fig. 6**). In some cancer types, such as HNSC, the co-deleted region is very small and a single driver gene, *CSMD1*, was identified (**Fig. 2f**). In other tumor types, such as READ and SARC (**Fig. 2g-h**), larger deleted regions that encompass *CSMD1* were identified, with *CSMD1* being the most commonly deleted gene in all cases. This strongly suggests that *CSMD1* – a known tumor suppressor in various cancer types[29], [30], [31] not linked before to Chr8p loss – is an important (albeit not necessarily sole) driver of this aneuploidy.

Overall, this analysis identified 96 potential driver genes, within 31% of the CA-CT pairs. Of these, in about 70% of the cases a single gene was nominated, whereas in the rest of the cases two or three drivers were proposed (**Fig. 2i** and **Supp. Table 2**). The analysis revealed new potential drivers of recurrent arm losses, such as *CSMD1* in Chr8p, in addition to strong TSGs that have been previously assumed to be aneuploidy drivers, such as *PTEN* in Chr10q.

### Point mutations co-occurring with arm losses

Next, we investigated genes that are frequently mutated when the arm on which they reside is lost (pattern PM+AL, **Fig. 3a**). For each CA-CT pair, we applied MutSig2CV [32], a tool designed to identify genes mutated more often than expected by chance given the background mutation processes in the dataset, to the group of samples harboring the arm loss, focusing on genes located on that arm. The results for BRCA are shown in **Fig. 3b** and for all other tumor types in **Supp. Fig. 7**. The full list of genes that were identified in this pattern analysis is provided in **Supp. Table 2**.

**Figure 3:**
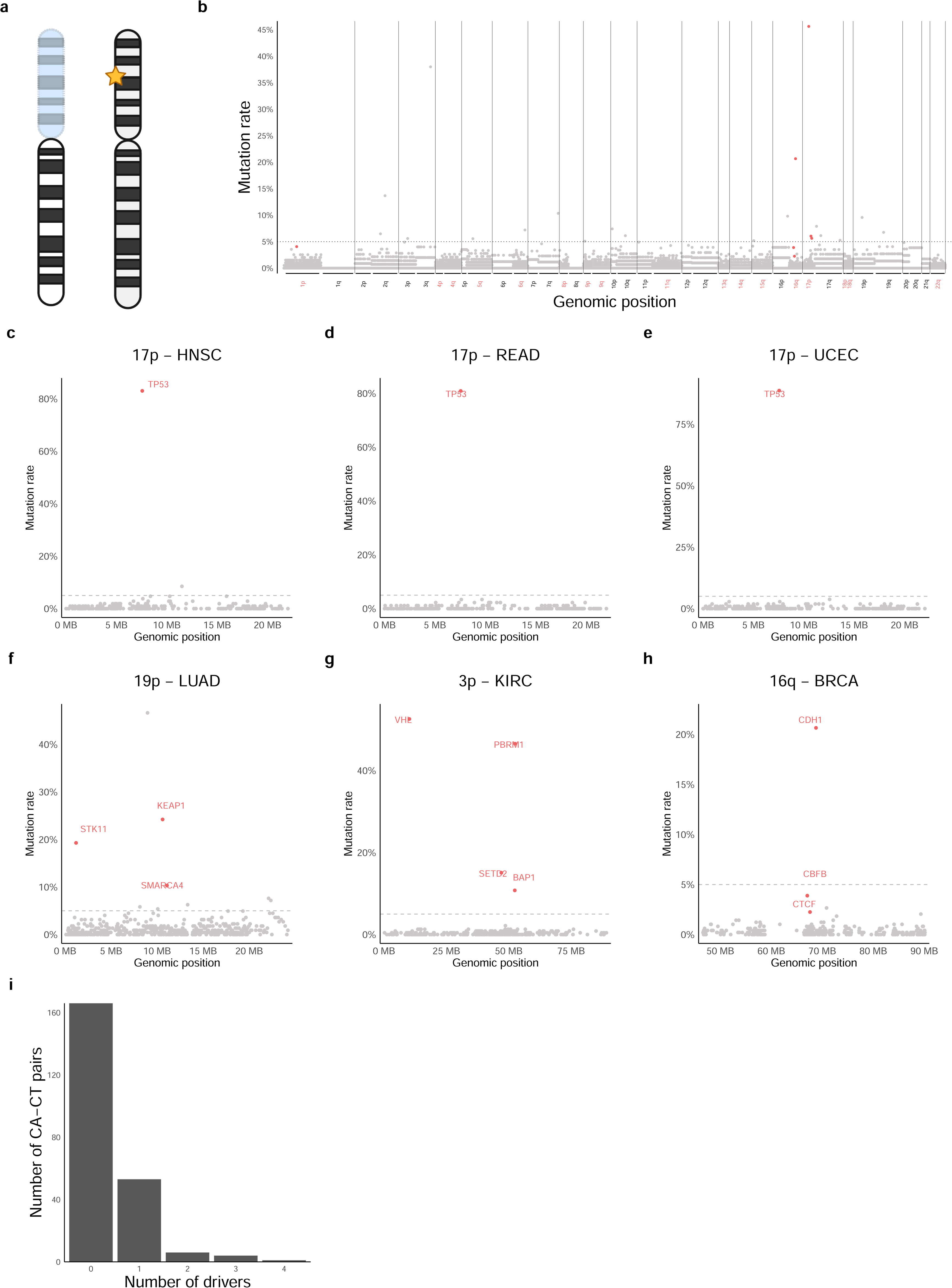
Point mutations co-occurring with arm losses. **a. Schematic representation of the PM+AL inactivation pattern.** **b. The prevalence of gene mutations co-occurring with arm losses in BRCA.** Mutation rate of each gene when the other copy of the arm on which it resides is lost in BRCA samples. Arms commonly deleted are colored in red. Genes identified as drivers by this pattern are colored in red. **c. + d. + e. The prevalence of gene mutations co-occurring with Chr17p loss in HNSC, READ and UCEC**. **f. The prevalence of gene mutations co-occurring with Chr19p loss in LUAD**. **g. The prevalence of gene mutations co-occurring with Chr3p loss in KIRC**. **h. The prevalence of gene mutations co-occurring with Chr16q loss in BRCA**. **i. Distribution of the number of drivers identified in CA-CT pairs for the PM+AL pattern.**

A notable example of this pattern is *TP53*, which is frequently mutated in conjunction with a Chr17p loss. This association was observed in virtually all cancer types with frequent loss of Chr17p (**Fig. 3c-e** and **Supp. Fig. 8**). *TP53* is a well-established driver of Chr17p loss, and its bi-allelic inactivation through a combination of mutation and arm loss was observed in several cancer types before [16], [17], [18].

Unlike *TP53*, most genes identified in this analysis are frequently mutated in only one or two cancer types. For example, while Chr19p loss is common across three cancer types, candidate driver genes fitting this pattern for this recurrent arm loss were identified only in LUAD (**Fig. 3f**). Similarly, for losses of Chr16q and Chr3p, genes fitting this pattern were found only in three and in six of the nine relevant cancer types, respectively. In both cases, the identified driver gene were cancer type-specific: for example, *TGFBR2* was nominated as a driver of Chr3p loss only in HNSC, whereas *VHL* and *SETD2* were nominated as drivers of the same aneuploidy only in KIRC (**Fig. 3g** and **Supp. Fig. 9**). Likewise, *CDH1* was nominated as a driver of Chr16q loss only in BRCA (**Fig. 3h** and **Supp. Fig. 9**). *CDH1* was also identified as a driver in UCEC through the analysis of FD-AL pattern (see below).

Overall, this analysis identified a total of 81 potential driver genes, within 28% of the CA-CT pairs. Of these, in the vast majority (83%) of cases, a single gene was identified, whereas in the other cases, the impacted region contained multiple (2 to 4) genes (**Fig. 3i** and **Supp. Table 2**). This analysis identified well-known TSGs, such as *TP53* in several tumor types, *VHL* [33] in kidney cancer, and *SMARCA4* [34] in lung cancer, but also genes that had not been previously implicated in driving recurrent arm losses, such as *NSD1* and *RASA1* in LUSC and HNSC, respectively (**Supp. Fig. 9**).

### Focal deletions mutually exclusive with arm losses

Some driver genes may be lost either through an arm loss or through a more focal loss, but not by both losses simultaneously. This is either because the driver itself is an essential gene that cannot be bi-allelically inactivated, or because of essential genes in close proximity to the driver. Therefore, we investigated focal deletions that are prevalent in a particular arm only when the other copy of the arm is not lost (pattern FD-AL, **Fig. 4a**). These mutually exclusive events were relatively large, spanning large fractions of the respective arm (a median length of 6.9Mb and of 160 genes; exemplified for BRCA in **Fig. 4b** and for all other tumor types in **Supp. Fig. 10**). For each CA-CT pair, we applied GISTIC2.0 [27] to samples lacking the arm loss to identify such focal losses. Since the identified chromosomal regions were large, harboring tens to hundreds of genes, most of which very likely to be mere passengers, extra steps of filtering were required for driver nomination. We therefore used PRODIGY [35], a tool that we previously developed to identify driver genes based on their proximity in a protein-protein interaction network to members of dysregulated pathways. To identify dysregulated pathways linked to the deletion of the region, we performed differential gene expression analysis by comparing samples with focal or arm loss of the region to all other samples. Using the hypergeometric test, we analyzed pathways from the Reactome pathway database [36] and identified those enriched with dysregulated genes. Next, we employed PRODIGY to rank the genes that reside within the region based on their association with these dysregulated pathways (see Methods). However, we noticed that many of the nominated candidates were in fact known oncogenes, likely due to the large number of genes per region and the lack of directionality requirements in our analysis. Therefore, to increase the confidence in the identified candidate genes, we further refined our approach to focus solely on known TSGs. While this filtering step limited the scope of this particular analysis, it greatly increases the confidence in the validity of the identified drivers.

**Figure 4:**
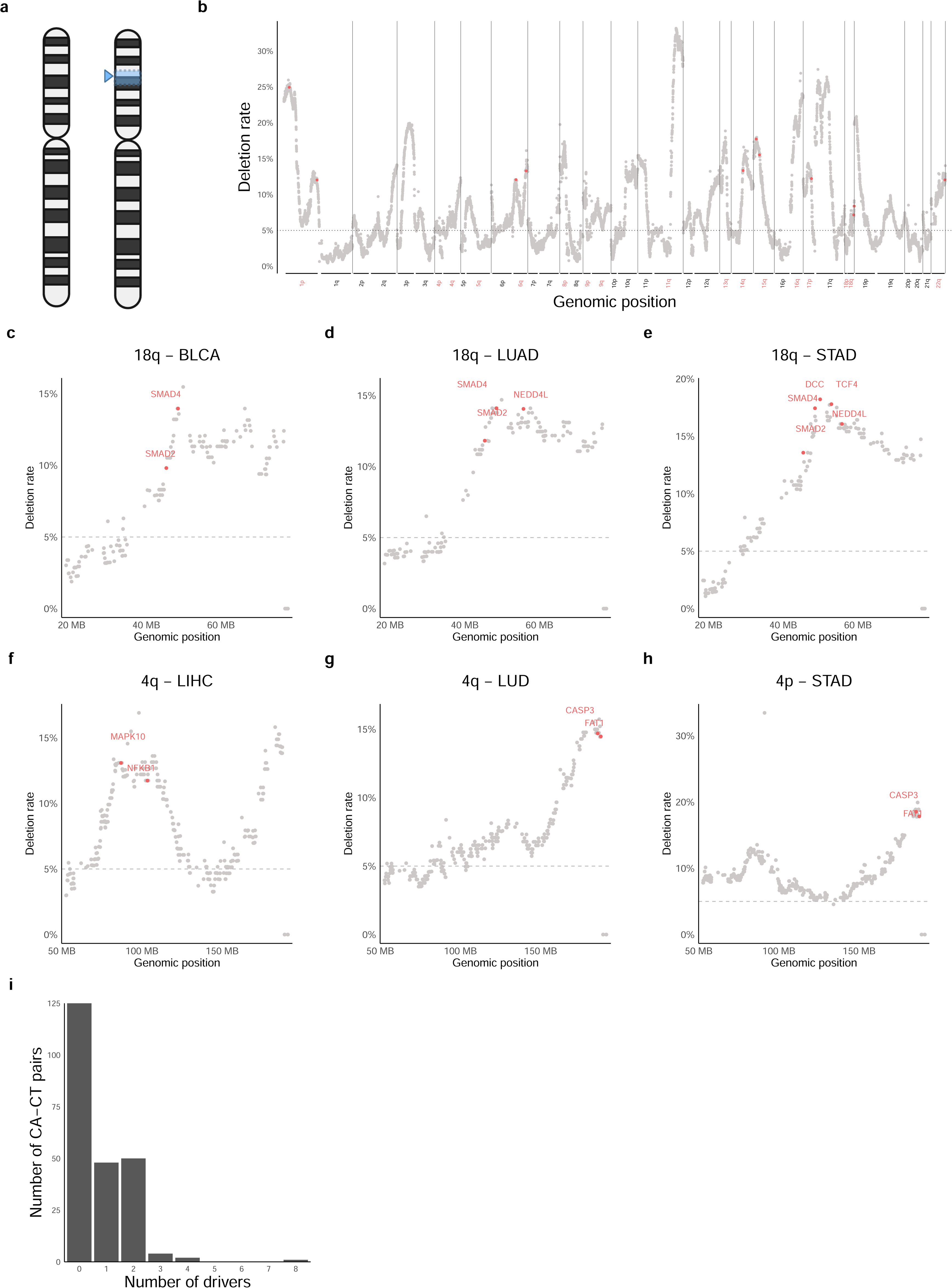
Focal deletions mutually exclusive with arm losses. **a. Schematic representation of the FD-AL inactivation pattern.** **b. The prevalence of gene-level focal deletions when the other copy of the arm is not lost in BRCA.** Focal deletion rate of each gene given that the other copy of the arm on which it resides is not lost in BRCA samples. Arms commonly deleted are colored in red. Genes identified as drivers by this pattern are colored in red. **c. + d. + e. The prevalence of gene-level focal deletions when Chr18q is not lost in BLCA, LUAD and STAD**. **f. + g. + h. The prevalence of gene-level focal deletions when Chr4q is not lost in LIHC, LUAD and STAD**. **i. Distribution of the number of drivers identified in CA-CT pairs for the FD-AL pattern.**

The full list of candidate drivers identified in this pattern analysis is provided in **Supp. Table 2**. An example of this pattern is seen on Chr18q, which contains a sizable region frequently lost in the absence of the arm loss across multiple cancer types. Our analysis identified *SMAD4* as a candidate driver of Chr18q loss in eight cancer types, while also nominating other genes as additional, cancer type-specific drivers (**Fig. 4c-e** and **Supp. Fig. 11**).

Another interesting example is that of Chr4q, for which different regions were found to be frequently lost, in a tumor type-dependent manner (**Fig. 4f-h** and **Supp. Fig. 12**). One region, located at the end of that arm, was observed across several cancer types, and it includes two prominent candidate driver genes, *CASP3* and *FAT1*. *CASP3* is a known TSG via its role in apoptosis [37], and *FAT1* is an atypical cadherin that was also shown to function as a TSG in some cancer types [38]. Another region, situated in the middle of Chr4q, appeared only in a subset of the cancer types, such as LIHC, suggesting that the drivers that reside within that region are tumor type-specific (**Fig. 4f** and **Supp. Fig. 12**).

Overall, this analysis identified 176 potential driver genes, within 46% of the CA-CT pairs (**Fig. 4i** and **Supp. Table 2**). Of these, in almost all cases (93%) one or two genes were identified. Some of these genes, like *CASP3* and *MAPK10*, have not been proposed as aneuploidy drivers before.

### Point mutations mutually exclusive with arm losses

Applying the same logic, we next searched for genes that are frequently mutated in the absence of arm-level loss. For each CA-CT pair, we identified genes that were mutated significantly more frequently when the other copy of the arm on which they reside was not lost (pattern PM-AL, **Fig. 5a**). We identified 374 candidate genes, all of which were found in the UCEC, STAD, and COAD-READ cancer types. However, in contrast to the patterns described above, the genes that were identified in this analysis were spread more or less evenly throughout the genome, with most genes presenting a surprisingly high mutation rate, as demonstrated in **Fig. 5b** for UCEC (See **Supp. Fig. 13** for the other tumor types). Furthermore, only ∼10% of these genes were known TSGs, in comparison to 68% and 83% in the FD+AL and PM+AL patterns, respectively. (The FD-AL pattern analysis considered only TSGs to begin with.)

**Figure 5:**
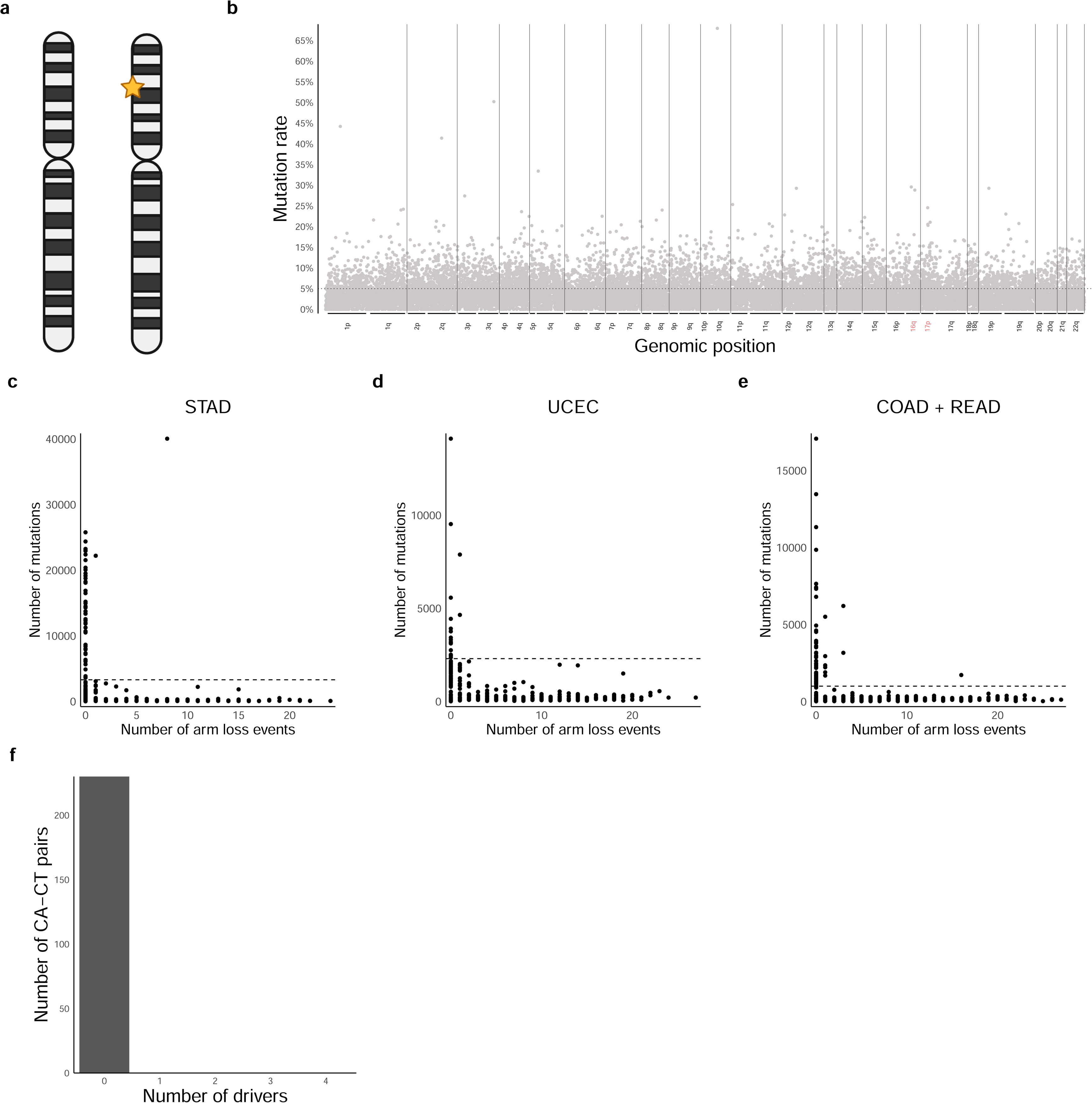
Point mutations mutually exclusive with arm losses. **a. Schematic representation of the PM-AL inactivation pattern.** **b. The prevalence of gene-level mutations in the absence of arm loss in UCEC.** Mutation rate of each gene when that the other copy of the arm on which it resides is not lost in UCEC samples. Names of arms commonly deleted are colored in red. **c. + d**. + **e. Relationship between the number of mutations and number of arm deletions in STAD, UCEC, COAD and READ samples.** Each dot shows the rates for one sample. **f. Distribution of the number of drivers identified in CA-CT pairs for the PM-AL pattern.**

These results suggest that the genes identified might not be true aneuploidy drivers. Importantly, the four cancer types in which the genes were identified are all known to have hyper-mutated, chromosomally stable subtypes [39], [40], [41]. Indeed, we found a subgroup of samples in these cancer types with a very high mutation count but very few, or even zero, arm losses (**Fig. 5c-e**), suggesting that the high mutation rate in these genes was mutually exclusive to arm losses simply because of this strong negative association, indicating that these were not true driver genes (**Fig. 5f**).

### Integrating the genetic analyses with the gene expression analyses

Overall, we identified 311 CA-CT candidate drivers involving 135 unique genes. Approximately 86% of these are known TSGs. Importantly, however, our analysis nominated only ∼7.6% of all known TSGs as putative drivers of arm losses, highlighting the value of harnessing genomic data to distinguish putative drivers from TSGs that just happen to reside on a recurrently lost arm.

Most of the candidate driver genes fit only one of the inactivation patterns discussed above (**Fig. 6a**), highlighting the importance of employing complementary approaches for their nomination. Genes that fit multiple patterns are particularly noteworthy, though, as they provide stronger evidence supporting their roles as aneuploidy drivers. *FAT1*, *PTEN*, *RB1*, and *SMAD4* are the only genes that fit all three patterns. Indeed, all except *FAT1* were previously suggested to drive their respective arm loss [19], [42], [43].

**Figure 6:**
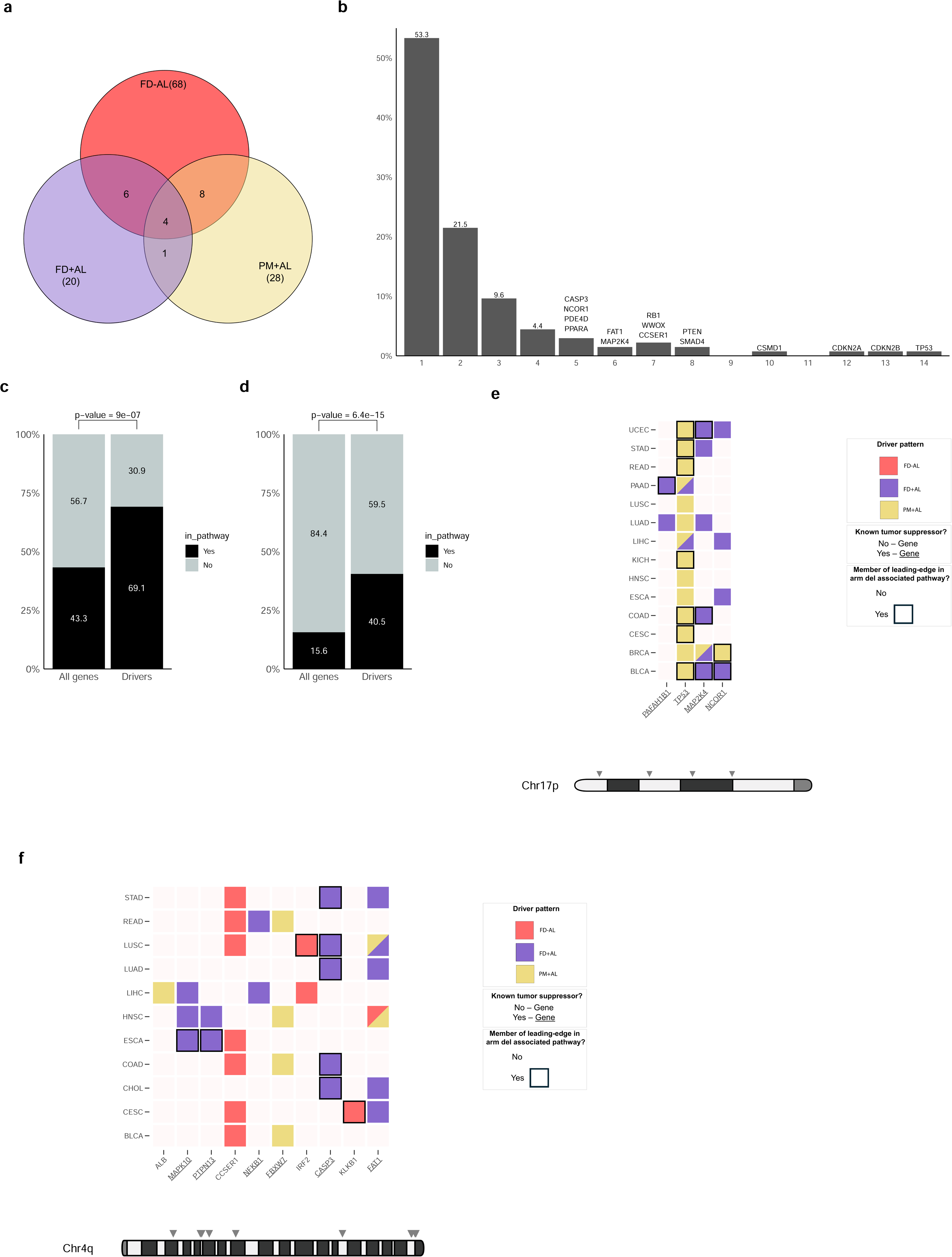
Integrating the genetic driver analysis with the gene expression analysis. **a. Venn diagram of the candidate drivers identified by each inactivation pattern.** **b. Recurrence of candidate drivers across cancer types.** The histogram shows the distribution of drivers according to the number of cancer types in which they were identified. Drivers that occurred in five or more cancer types are named above their bar. **c. Enrichment of drivers in aneuploidy-associated pathways.** For each CA-CT pair, the dysregulated Reactome pathways were identified by performing GSEA between the expression in tumor samples with the arm loss and those without it. The plot shows the fraction of drivers vs. the fraction of all genes belonging to such pathways. P-value is calculated using Fisher’s Exact test. **d. Enrichment of drivers in the leading-edge of aneuploidy-associated pathways.** The plots are as in c but showing the fraction of drivers vs. the fraction of all genes belonging to the leading-edge subsets of the dysregulated pathways. **e. A summary visualization of the candidate drivers of Chr17p loss across cancer types.** The candidate drivers of Chr17p loss identified in each cancer type are shown in a matrix with rows corresponding to the cancer types and columns corresponding to the drivers. The colors reflect the perturbation pattern(s) in which each gene was identified. Drivers present in the leading-edge subsets of at least one dysregulated pathway are highlighted with a black border. The names of known TSGs are underlined. The chromosome diagram shows the locations of the drivers, in the order they appear in the matrix. **f. A summary visualization of the candidate drivers of Chr4q loss across cancer types.** Same plot as in e, but for Chr4q.

Interestingly, most putative drivers are specific to only one or a few cancer types (**Fig. 6b**). However, a subset of genes appears in multiple cancer types, and 15 genes (11% of the proposed drivers) were identified in five or more cancer types, highlighting their potential significance as universal drivers (**Fig. 6b**). For example, *TP53*, the most common TSG in human cancer, was identified as a driver of Chr17p loss in 14 different tumor types.

To further validate our findings and highlight the most interesting candidate drivers, we assessed whether the putative drivers were enriched for genes that belong to pathways that are significantly altered upon that arm loss. To do so, we applied GSEA to the gene expression results from the within-cancer type comparisons discussed above (**Fig. 1**) using the 1,293 pathways of size between 10 and 500 from the Reactome pathway database [36]. We identified 33,412 such dysregulated pathways across all 230 CA-CT pairs. We found that our putative drivers were indeed significantly more likely to participate in dysregulated pathways (**Fig. 6c**).

Additionally, we observed that the putative drivers were highly enriched in the leading-edge subsets of the dysregulated pathways, defined as the subsets of genes with the highest contribution to a pathway’s enrichment signal (**Fig. 6d**; see Methods).

These results increase our confidence that pathways containing drivers in their leading-edge are more likely to be directly associated with the respective arm loss. Similarly, they boost our confidence that our approach indeed identifies biologically meaningful driver genes, which affect the gene expression patterns of the tumors. We therefore integrated the genetic analyses with the gene expression analyses. For example, in tumor types in which Chr17p is commonly lost, the cell cycle checkpoint and programmed cell death pathways are downregulated in comparison to Chr17p-wildtype samples across several cancer types (**Supp. Table 1**). *TP53* appears in the leading-edge of these gene sets, and is proposed by our analysis to be a strong driver of this common aneuploidy (**Supp. Table 2**). Similarly, Chr4q loss is associated with the downregulation of cell death pathways in STAD, COAD, and LUSC (**Supp. Table 1**), presumably driven by the inactivation of *CASP3* (**Supp. Table 2**). Another noteworthy observation, seen in BLCA, GBM, and HNSC, is that the apoptosis, cell cycle checkpoint, and programmed cell death pathways are upregulated in tumors with Chr9p loss in comparison to healthy tissues, but are downregulated in comparison to Chr9p-wildtype samples (**Supp. Table 1**). This pattern suggests that these pathways may be generally upregulated in certain cancer types, with Chr9p loss mitigating this detrimental activation through the downregulation of *CDKN2A* and *CDKN2B*, which we identify as putative drivers of this aneuploidy (**Supp. Table 2**).

Lastly, we provide a visual summary of the putative drivers of each CA-CT pair (**Fig. 6e,f** and **Supp. Fig. 14**). For instance, **Fig. 6e** provides an integrated view of the putative drivers of Chr17p loss. *TP53* appears across all cancer types, primarily through the PM+AL pattern. Three other genes are known TSGs that were proposed as drivers of this arm loss in different cancer types. These genes also frequently appear in the leading-edge subsets of dysregulated pathways identified in the GSEA analysis (**Fig. 6e**). In contrast, the putative drivers of Chr4q loss reveal a more diverse, tumor-type-specific group of drivers, including both known TSGs and less well-characterized genes (**Fig. 6f**). Most of these proposed drivers are specific to a handful of cancer types. Notable candidates include *CCSER1*, residing in a small region frequently bi-allelically deleted across many cancers; *CASP3*, a known TSG frequently focally deleted in the absence of Chr4q loss; and *FAT1*, another well-established TSG identified across many cancer types through all three patterns. Summary plots for all other recurrently lost arms are shown in **Supp. Fig. 14**. The full lists of putative drivers, together with the patterns and the pathways to which they belong, are provided in **Supp. Table 2**.

## Discussion

In this study, we presented a framework for identifying driver genes associated with cancer-type-specific recurrent arm losses. Our analysis revealed that focal deletions co-occurring with losses of the same arm are typically small (median size of ∼350Kb). The co-occurrence of these events suggests the presence of a genetic element that is bi-allelically inactivated by these deletions, thereby driving their recurrence. This observation enabled us to narrow down our search to specific regions of the arm and, in many cases, to propose candidate genes that likely contribute to the frequent loss of the arm.

Conversely, our analysis also identified medium-size deletions (median size of ∼6.9Mb) that are prevalent only in the absence of an arm loss. This pattern suggests the presence of a genetic element that is frequently inactivated by either event, driving the recurrence of these losses.

This pattern may suggest that only a mono-allelic inactivation of this driver is advantageous for the cancer cell. Alternatively, it may imply the existence of essential genes that reside in the close proximity of the driver gene(s), thereby reducing the likelihood of that region to be bi-allelically lost. Our results from the point-mutation analysis lends support to the latter explanation, as we could not identify a driver gene that was recalcitrant to bi-allelic inactivation.

Analysis of mutation data identified a class of genes frequently bi-allelically inactivated by an arm loss and a point mutation. *TP53* is the best-known example of this mechanism, and we indeed identified it as a strong driver of Chr17p loss across many tumor types. On the other hand, we see no evidence for the existence of genes whose inactivation by an arm loss and by a gene mutation is mutually exclusive, suggesting that essential genes whose complete loss is not compatible with cell fitness are rarely (if ever) drivers of arm losses. These results suggest that negative selection does not act against the bi-allelic inactivation of driver genes. Instead, the negative selection observed in FD-AL pattern appears to target the bi-allelic inactivation of nearby passenger genes rather than the driver gene itself.

By comparing the gene expression of cancer samples harboring a specific arm loss to those without it, followed by a gene set enrichment analysis, we identified pathways associated with the arm loss. However, our analysis demonstrates that these gene expression changes are not sufficient in and of themselves for driver identification, as they may be confounded by co-occurring aneuploidies and other genomic alterations. Therefore, an integrative approach that takes into account both gene expression and genetics, like the one we present here, is required. Integrating the genetic analysis with the gene expression analysis resulted in the identification of specific pathways whose dysregulation is most likely attributed to the arm loss. This approach also allowed us to link specific driver genes to these pathway alterations, pinpointing specific cellular pathway(s) through which the loss of these genes promotes cancer.

We note that most of our proposed candidates are cancer-type-specific, emphasizing the context specific role of arm loss (reviewed in [1]). A small subset of genes appears across cancer types, though, alluding to their strong and global role as cancer aneuploidy drivers. Additionally, the vast majority of drivers are recognized in only one of our analyzed genomic patterns, indicating that specific genes tend to prefer a specific mechanism of inactivation, and emphasizing the importance of such integrative analyses.

Our work successfully identified driver genes underlying common arm losses. However, there are several limitations to our approach: (1) We only focused here on arm losses; future research should expand this approach to include arm gains as well. (2) We only considered the most common focal gene inactivation mechanisms, namely mutations and copy number alterations. Incorporating a broader range of inactivation mechanisms (e.g., promoter methylation) into this framework may help capture more diverse driver patterns. (3) Our analysis considered only one event at a time, ignoring the potential driving role co-occurring aneuploidies [7], [9]. Much more data is needed in order to perform such combinatorial analysis, but with the fast accumulation of genomic information, this will likely become possible within a few years. (4) Finally, we only considered protein-coding genes. It will be important to ultimately extend the analysis to consider other genetic elements, such as microRNAs and long non-coding RNAs. Despite all of these limitations, we believe that this study makes an important step toward a systematic, comprehensive characterization of the drivers of aneuploidy in human cancer.

## Methods

### Data collection and processing

Data from 7,503 TCGA samples spanning 20 cancer types were obtained as follows: Gene expression and mutation data from data release version 35 were downloaded using the R package TCGAbiolinks [44], while segmentation (.SEG) files from TCGA data version 2016_01_28 were obtained from the Broad Institute’s FireBrowse website. Arm-level status for each sample and aneuploidy scores (defined as the total number of arm copy-number alterations) were extracted from Supplementary Table S2 of Taylor et al. [2]. A list of 1414 known tumor suppressor genes was compiled from the Cancer Gene Census [45] and the Tumor suppressor gene database v2.0 [46]. An arm-loss event was defined as common in a specific cancer type if it occurred in 20% or more of the samples.

### Differential gene expression analysis

Two types of differential gene expression analyses were performed to evaluate the impact of all recurrent arm losses on gene expression within each cancer type. The first analysis compared cancer samples with the arm loss to normal samples, using DESeq2 [47]. The second analysis compared cancer samples with the arm loss to those without it, by estimating the log fold-change of each gene, while accounting for differences in aneuploidy levels between the two groups. Since the sample’s aneuploidy level and the arm deletion status are correlated, inverse probability of treatment weighting (IPTW) [48] was used to account for the effect of the sample’s aneuploidy levels. Weights were calculated using the samples’ aneuploidy scores: first, the probability of a sample to harbor an arm loss, *P*(*has arm loss*), was estimated from the data. Then, a logistic regression model *L*(*p*|*s*), estimating the probability *p* of a sample to have the arm loss given its aneuploidy score *s*, was trained. Finally, the sample’s weight was calculated by dividing the observed probability of a sample to harbor the arm loss by the estimated probability of harboring the arm loss event given the sample’s aneuploidy score: 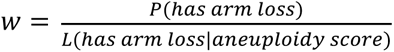. For each gene the log fold-change in gene expression between the two groups was estimated using a weighted version of Cohen’s d statistic, applied to *log*_10_ (*FPKM*) expression values and using the calculated sample weights: 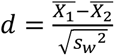, where *X̅* is the weighted mean of each group and *s*_*w*_^2^ is the weighted pooled standard deviation.

Each type of differential gene expression analysis was followed by GSEA [22], using the R clusterProfiler package [49], applied to the list of all differentially expressed genes using the “GSEA” function with default parameters. Two collections of gene sets were queried in each analysis: the MSigDB Hallmark gene sets (50 sets) [23] and the Reactome database [36], containing 1615 gene sets (only the1293 gene sets with 10 ≤ *size* ≤ 500 were used). For the analysis presented in **Fig. 1** and **Suppl. Fig. 1 and 2**, which incorporate results from multiple GSEA runs, p-values were first corrected across all results using the Benjamini-Hochberg procedure and then only results with q-value < 0.05 were taken.

### Analysis of focal deletions co-occurring with arm losses (FD+AL)

For each cancer type in which a particular arm loss event was common, we focused on the subset of tumors harboring the arm loss. Chromosomal regions in that arm affected by focal deletions co-occurring with the arm loss were identified by applying GISTIC2.0 [27] with the parameters used in [50]. Out of the genes within each identified region we excluded those deleted in <5 samples or in <5% of the samples with the respective arm loss. In regions containing ≤3 genes after filtering, genes with the highest deletion rate were nominated as putative drivers. In regions with >3 genes, a gene was nominated as a putative driver only if it was also identified as a driver independently in another cancer type.

### Analysis of point mutations co-occurring with arm losses (PM+AL)

For each cancer type in which a particular arm loss event was common, we focused on the subset of tumors harboring the arm loss. Genes affected by point mutations co-occurring with the arm loss were identified by applying MutSig2CV [32] using a q-value threshold of 0.25. Out of the results only genes located on the corresponding arm were taken.

### Analysis of focal deletions mutually exclusive to arm losses (FD-AL)

For each cancer type in which a particular arm loss event was common, we focused on the subset of tumors lacking the arm loss. We identified chromosomal regions that are frequently lost only when the other copy of the arm they belong to is not lost, by applying GISTIC2.0 [27] with the parameters used in [50]. To focus only on focal loss events that are mutually exclusive to the arm loss, regions that have ≥50% overlap with a region identified in the focal deletions co-occurrence analysis above were excluded. Frequent losses spanning both arms or ≥90% of the arm were excluded as well.

To identify the most plausible drivers in each region, we adapted PRODIGY [35], a tool originally designed for patient-specific ranking of cancer driving mutations, to pinpoint drivers of large focal deletions. This tool ranks candidate genes by estimating their cumulative effect on dysregulated pathways, as identified by differential expression analysis. Specifically, given a protein-protein interaction network *G* = (*V*, *E*, *W*), where *W* are edge weights representing interaction confidence, a set of differentially expressed genes *DEG* and a list of dysregulated pathways for a pathway *p* represented by the graph *G*_*p*_ = (*V*_*p*_, *E*_*p*_) the effect of each candidate gene *g* is estimated as follows:

1. Constructing a new graph *G*_*p*,*g*_ = (*V*_*p*,*g*_, *E*_*p*,*g*_, *W*_*p*,*g*_, *P*_*p*,*g*_), where:

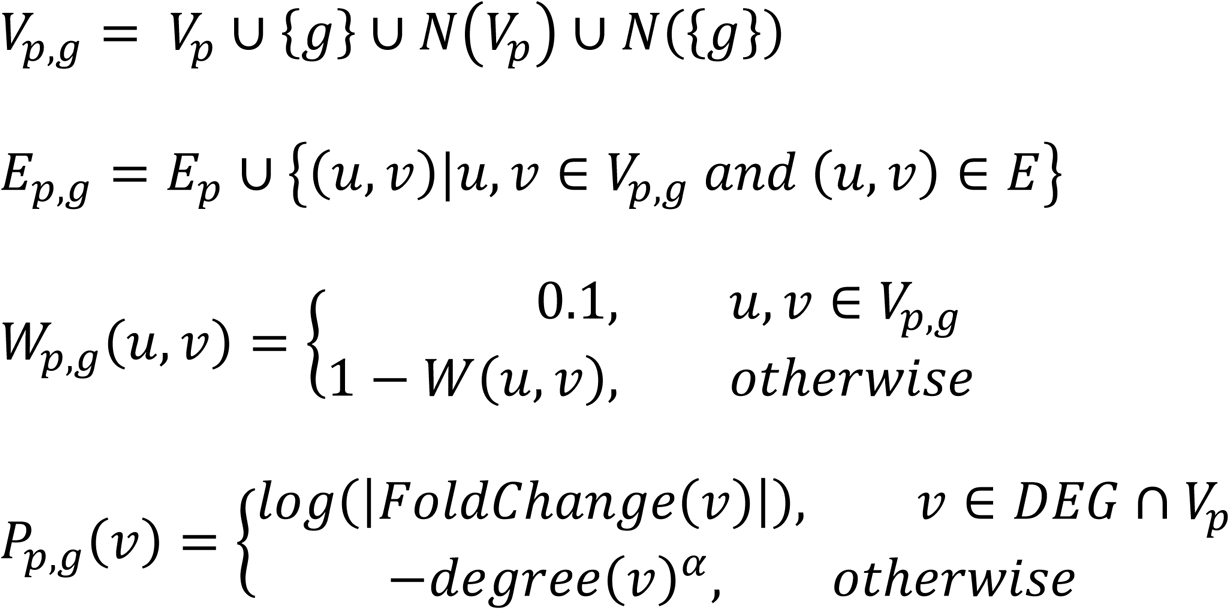

Here *N*(*S*) is the list of direct neighbors of set of vertices *S* in *G*and *⍺* is a parameter that controls the penalty assigned to nodes with high degree. A value of *⍺* = 0.05 was used, following the recommendation in the original paper.
2. Finding an approximate solution to the rooted prize collecting Steiner tree problem [51], i.e. a sub-tree *G*‵ = (*V*‵, *E*‵) rooted at *g* maximizing the score:

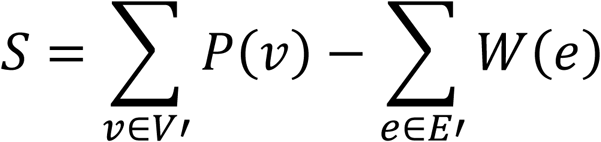
3. Assigning a normalized effect score by dividing the score *S* of the solution by the maximal possible score *S*_*max*_ = ∑_*v*∈*DEG*∩*Vp*′_ *log*(|*FoldChange*(*v*)|)

The following steps were applied to each chromosomal region:

1. First, differential expression analysis was conducted, comparing cancer samples with a focal or arm copy-number loss of the region to all other samples.
2. Second, pathways from the Reactome pathway database [36] enriched for differentially expressed genes were identified by the hypergeometric test.
3. Third, PRODIGY was used to rank the genes in the region by their cumulative effect on the dysregulated pathways, using the STRING protein–protein interaction network [52].
4. Known TSGs appearing in the top 5 ranking genes were proposed as putative drivers.

### Analysis of point mutations mutually exclusive to arm losses

For each cancer type in which a particular arm loss event was common, genes on that arm affected by point mutations at a significantly lower rate in tumors in which the other copy of the arm was lost, in comparison to tumors in which it was not lost, were identified by applying a Fisher’s Exact test. The p-values from all tests for each CA-CT pair were corrected using the Benjamini-Hochberg procedure and then filtered with a q-value threshold of 0.05. While a more precise version of this test would account for the reduced number of alleles for genes in tumors harboring the arm loss, no notable results were identified even with this less stringent approach.

### Enrichment analysis of driver genes in dysregulated pathways

For each CA-CT pair, differentially expressed genes between the cancer samples with the arm loss and without it were computed as described above, followed by GSEA using 1,293 pathways from the Reactome database. This analysis generated a collection *Θ* of 33,412 PT-CA-CT triplets, with each triplet representing a pathway (PT) that is dysregulated when comparing samples with a particular chromosome arm (CA) loss to those without the loss in a specific cancer type (CT). This collection was utilized to create a gene-level list *Ω* consisting of 1,060,326 GN-CA-CT triplets, where a triplet GN-CA-CT with gene GN belongs to *Ω* if GN belongs to a pathway PT for some triplet PT-CA-CT in *Θ*. Similarly, a list Δ of 186 DR-CA-CT triplets was created, where DR-CA-CT belongs to Δ if DR was nominated as a driver of that CA-CT pair in our prior analysis and DR-CT-CT belongs to *Ω*.

As a background set *Ω* ‵ for *Ω*, we used all possible triplets. Specifically, *Ω* ‵ consists of all GN-CA-CT such that GN is one of the 10,646 genes appearing in at least one of the 1293 pathways from the Reactome database, and CA-CT is one of the 230 CA-CT pairs. This set contained 2,448,580 triplets. Similarly, a background set Δ‵ for Δ was created, where DR-CA-CT belongs to Δ‵ if DR was nominated as a driver of that CA-CT pair in our prior analysis and DR-CA-CT belongs to *Ω*‵ (Δ‵ does not contain all 311 identified DR-CA-CT since not all drivers appear in the Reactome database).

Fisher’s exact test was applied to test for enrichment of drivers within the dysregulated pathways, comparing *Ω* and Δ to the background set. An analogous procedure was performed to test for enrichment of drivers within the leading-edge subsets of the GSEA results.

## Supplementary Figure Legends

**Supplementary Figure 1: Distribution of the number of altered ‘Hallmark’ pathways associated with recurrent chromosome-arm losses.** (a) Tumors with vs. without the chromosome-arm loss. (b) Tumors with the chromosome-arm loss vs. normal tissues.

**Supplementary Figure 2: Pathway dysregulation associated with recurrent chromosome-arm losses for each cancer type.** Each panel shows one cancer type. For each recurrent chromosome-arm loss in each cancer type, the upregulation or downregulation of cancer-related ‘Hallmark’ gene sets are shown, comparing tumors with vs. without each chromosome-arm loss. NES: GSEA normalized enrichment score.

**Supplementary Figure 3: Pathway dysregulation associated with recurrent chromosome-arm losses for each cancer type.** Same as **Supplementary** Figure 2 but comparing tumors with the chromosome-arm loss to normal tissues.

**Supplementary Figure 4: The prevalence of focal deletions co-occurring with arm losses in each cancer type.** Each panel shows one cancer type. Shown is the focal deletion rate of each gene when the other copy of the arm on which it resides is lost in the relevant cancer type. Commonly deleted chromosome-arms are colored in red. Genes identified as drivers that fit this pattern are colored in red.

**Supplementary Figure 5: The prevalence of focal deletions co-occurring with Chr10q loss in BLCA and CESC.**

**Supplementary Figure 6: The prevalence of focal deletions cooccurring with Chr8p loss in BLCA, BRCA, COAD, ESCA, KIRC, LIHC and LUSC.**

**Supplementary Figure 7: The prevalence of gene mutations co-occurring with chromosome-arm losses in each cancer type.** Each panel shows one cancer type. Shown is the mutation rate of each gene when the other copy of the arm on which it resides is lost in the relevant cancer type. Commonly deleted chromosome-arms are colored in red. Genes identified as drivers that fit this pattern are colored in red.

**Supplementary Figure 8: The prevalence of gene mutations co-occurring with Chr17p loss in BRCA, COAD, KICH, LIHC, LUSC, PAAD, STAD, CESC, ESCA, LUAD and BLCA.**

**Supplementary Figure 9: The prevalence of gene mutations co-occurring with Chr16q loss in LIHC and UCEC and with 3p loss in CHOL and HNSC.**

**Supplementary Figure 10: The prevalence of gene-level focal deletions when the other copy of the arm is not lost in each cancer type.** Focal deletion rate of each gene given that the other copy of the arm on which it resides is not lost in the relevant cancer type. Commonly deleted chromosome-arms are colored in red. Genes identified as drivers that fit this pattern are colored in red.

**Supplementary Figure 11: The prevalence of gene-level focal deletions when Chr18q is not lost in COAD, ESCA, READ, BRCA and LUSC.**

**Supplementary Figure 12: The prevalence of gene-level focal deletions when Chr4q is not lost in ESCA, HNSC, CHOL, COAD, READ, CESC and LUSC.**

**Supplementary Figure 13: The prevalence of gene-level mutations in the absence of chromosome-arm loss in STAD, COAD and READ.** Shown is the mutation rate of each gene when that the other copy of the arm on which it resides is not lost in the relevant cancer type. Commonly deleted chromosome-arms are colored in red.

**Supplementary Figure 14: A summary visualization of the candidate drivers of each chromosome-arm loss across cancer types.** The candidate drivers of all recurrent chromosome-arms losses are shown for each cancer type, with rows corresponding to the cancer types and columns corresponding to the identified driver genes. The colors reflect the perturbation pattern(s) that each gene belongs to. Drivers present in the leading-edge subsets of at least one dysregulated pathway are highlighted with a black border. The names of known TSGs are underlined. The chromosome diagram shows the locations of the drivers, in the order of their appearance in the matrix.

## Supplementary Table Legends

**Supplementary Table 1: Gene set enrichment analysis (GSEA) of differentially expressed genes.** Shown are ‘Hallmarks’ and ‘Reactome’ gene sets, for comparisons of tumors with vs. tumors without each recurrent chromosome-arm loss, and tumors with each chromosome-arm loss vs. the respective normal tissue.

**Supplementary Table 2: The full list of candidate driver genes.** Shown are the driver genes of each recurrent chromosome-arm loss, together with the inactivation pattern that each of them fits into, and the gene expression pathways that are significantly dysregulated upon that chromosome-arm loss and which their leading-edge subset includes that gene.

## Acknowledgments

The authors would like to thank all members of the Ben-David lab and the Shamir lab for their invaluable help and support throughout this study.

## Authors’ contributions

All authors conceived the study, designed the analyses, interpreted the results, and wrote the manuscript. R. Saad, performed the analyses. R. Shamir. and U. Ben-David. oversaw the study.

## Funding

Work supported by the European Research Council Starting Grant (grant #945674 to U. Ben-David), the Israel Science Foundation (grant #1805/21 to U. Ben-David and #2206/22 to R. Shamir), the Israeli Cancer Research Fund (Project Grant to U. Ben-David), and the BSF (Project Grant #2023236 to U. Ben-David). R. Saad was supported in part by a fellowship from the Edmond J. Safra Center for Bioinformatics at Tel Aviv University.

## Data availability

The code for all the analyses is available on GitHub at https://github.com/Shamir-Lab/aneuploidy-drivers-detection. The TCGA datasets that were analyzed were obtained using the TCGAbiolinks R package.

## Declarations

## Ethics approval and consent to participate

Ethics approval is not applicable

## Competing interests

U.B.-D. declares receiving consulting fees from Accent Therapeutics. R. Sa. Is a current employee of CytoReason LTD.

