## Supplementary Figures for "Integrating gene expression, mutation and copy number data to identify driver genes of recurrent chromosome-arm losses"

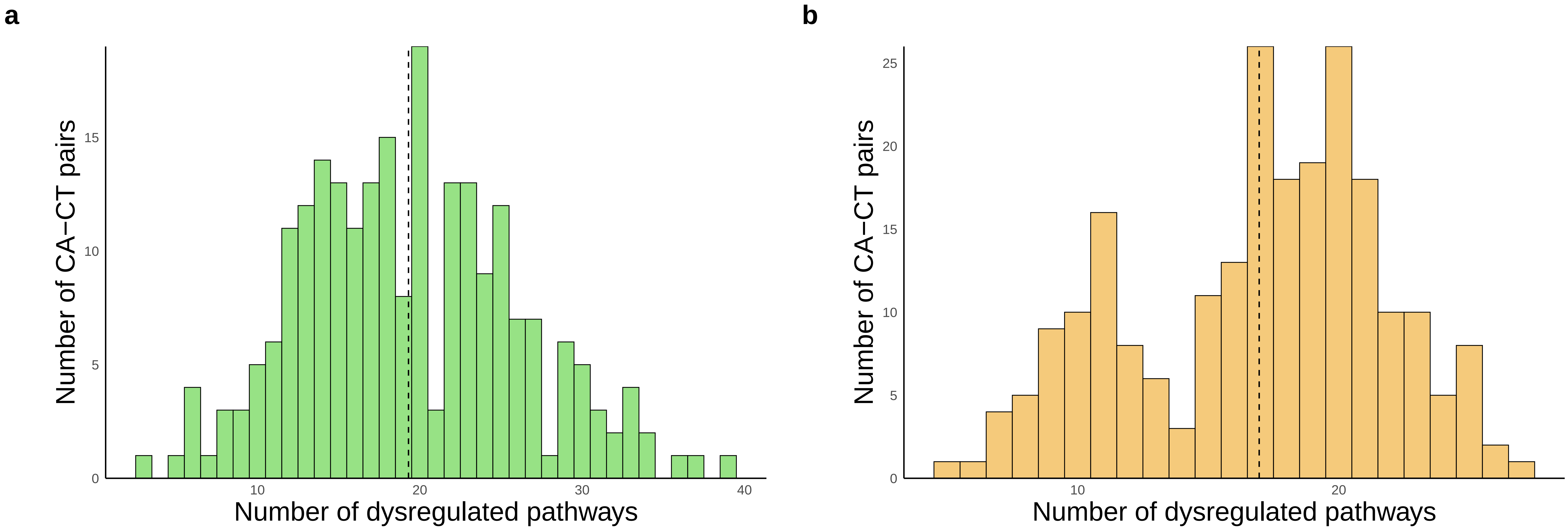


Supplementary Figure 1

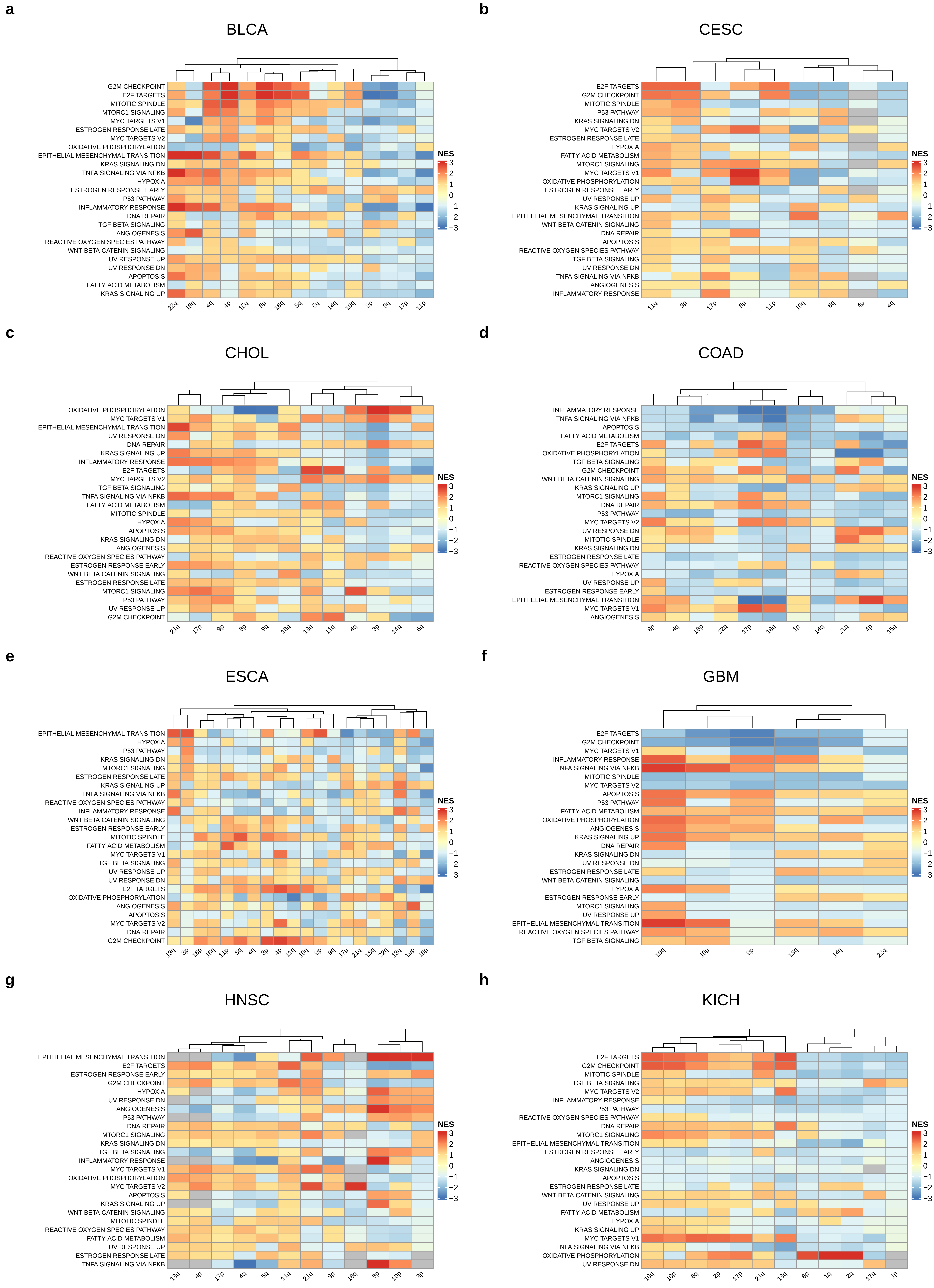


Supplementary Figure 2

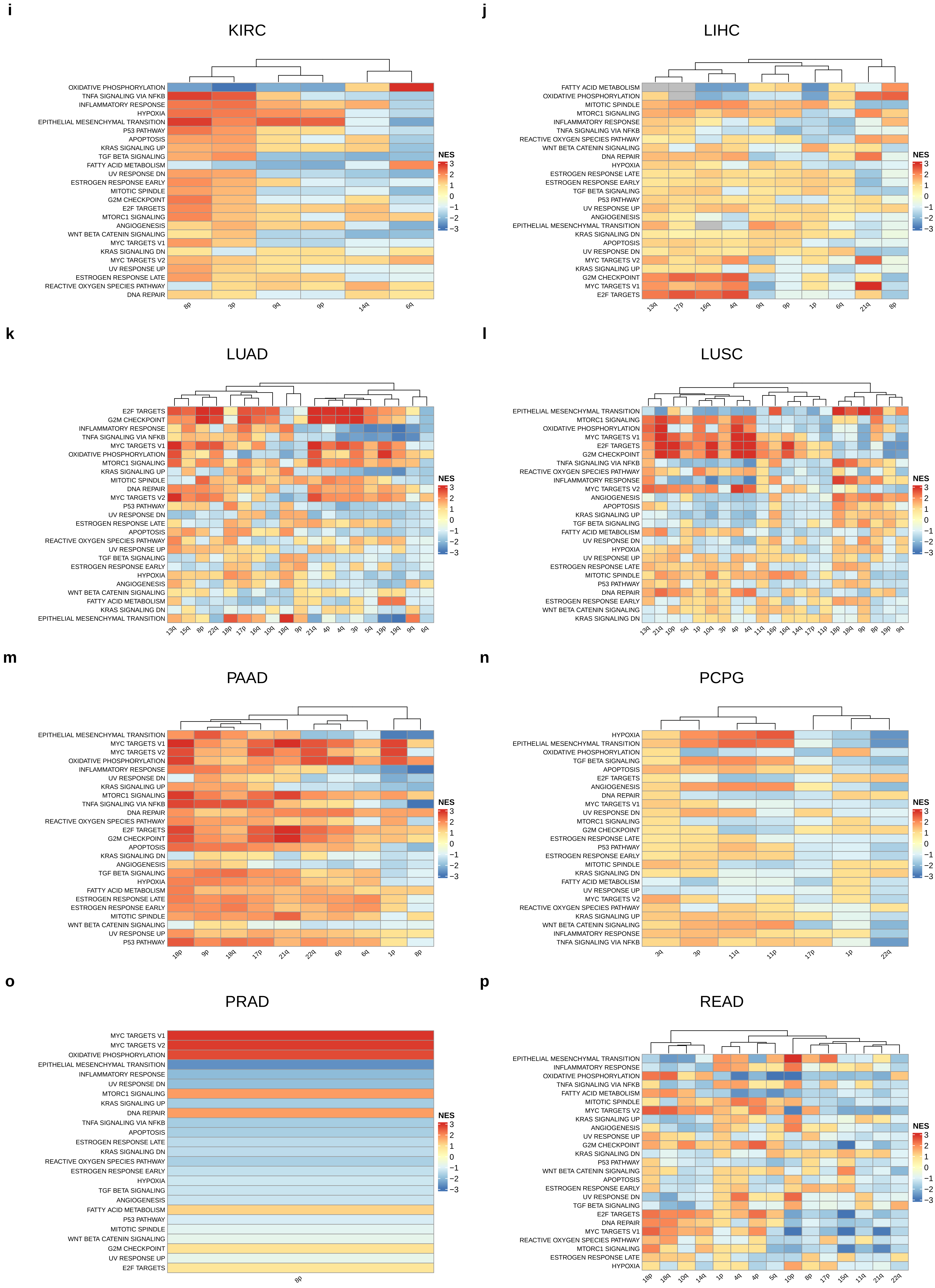


Supplementary Figure 2

**q**

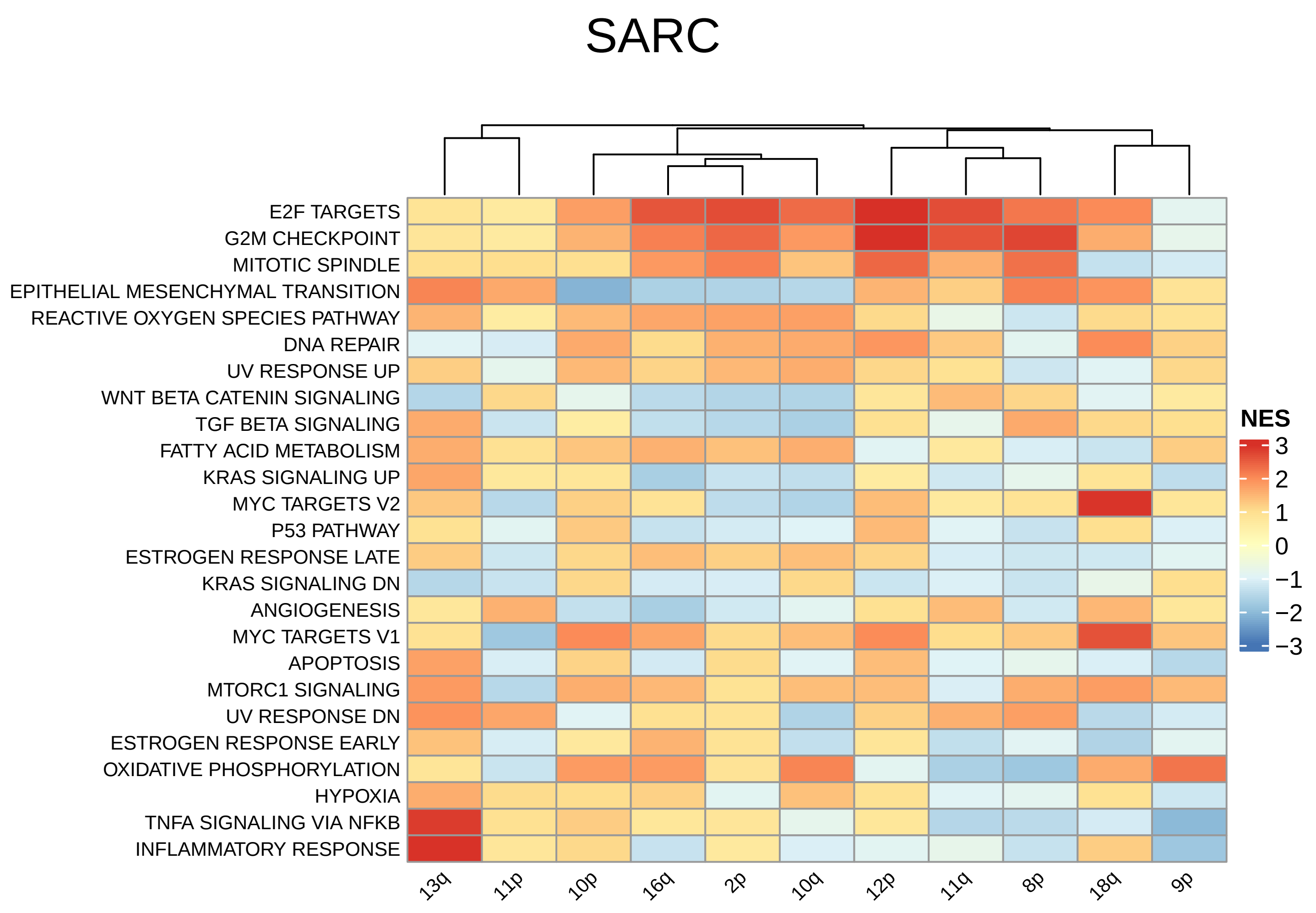


**r**

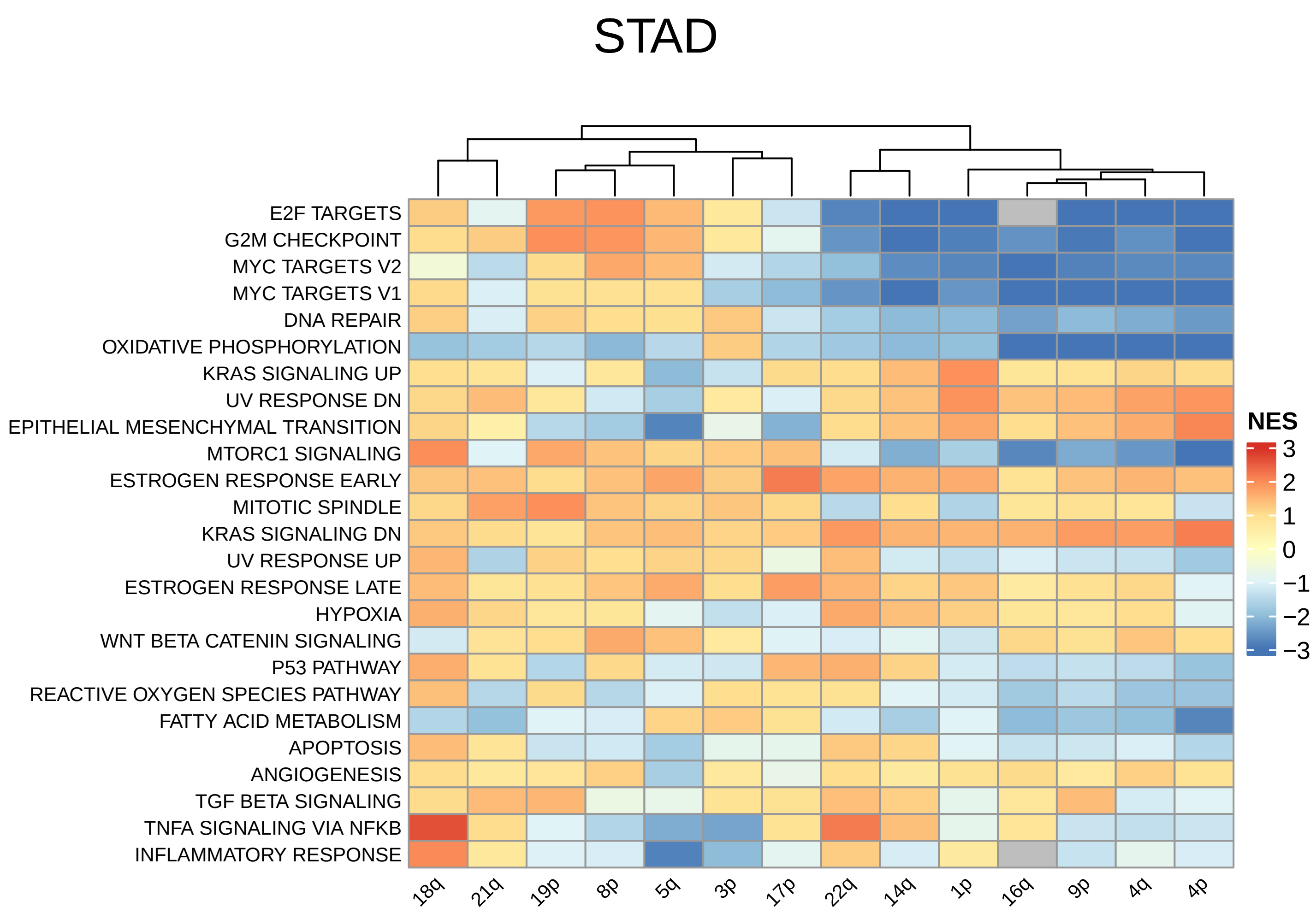


**s**

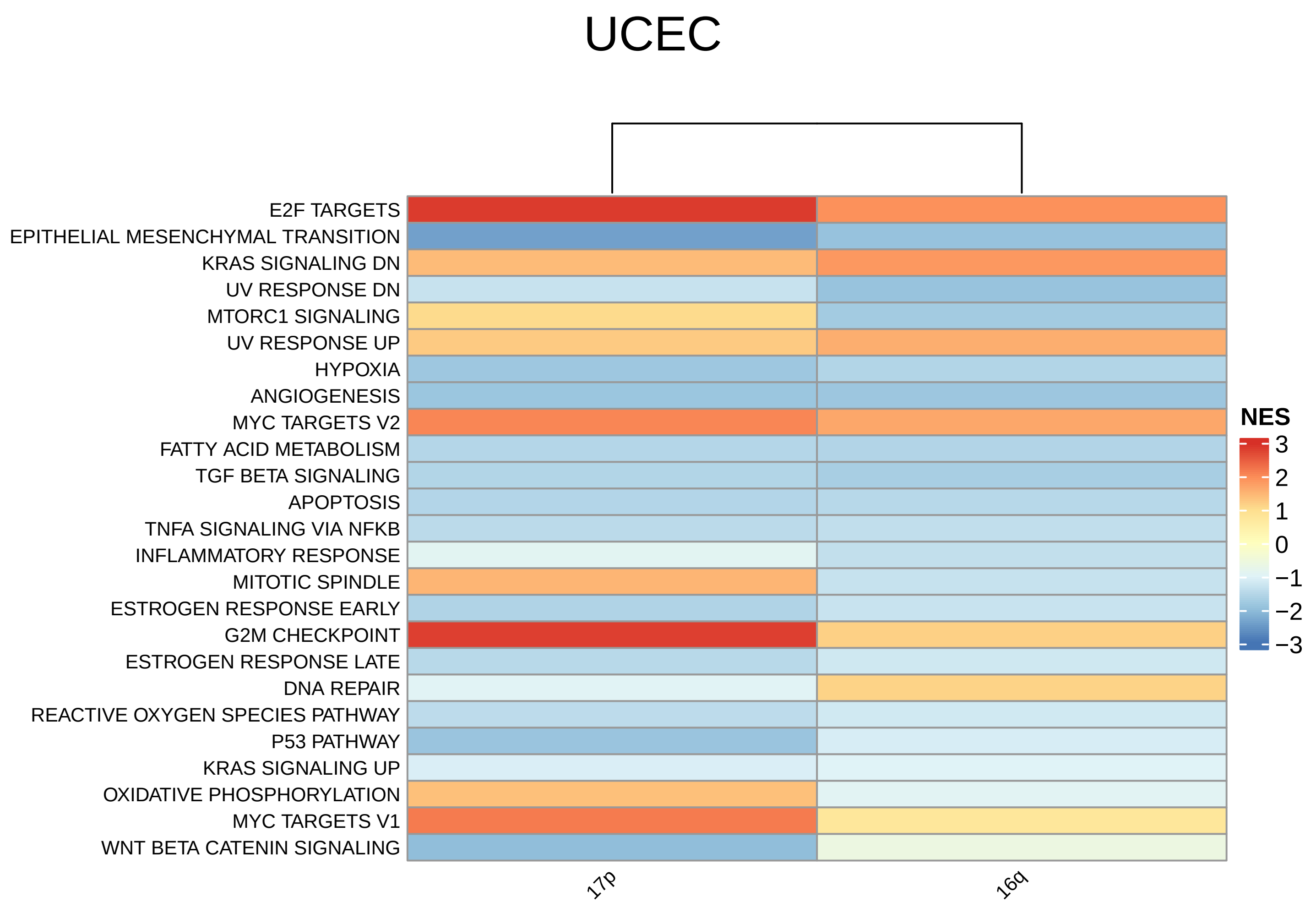


Supplementary Figure 2

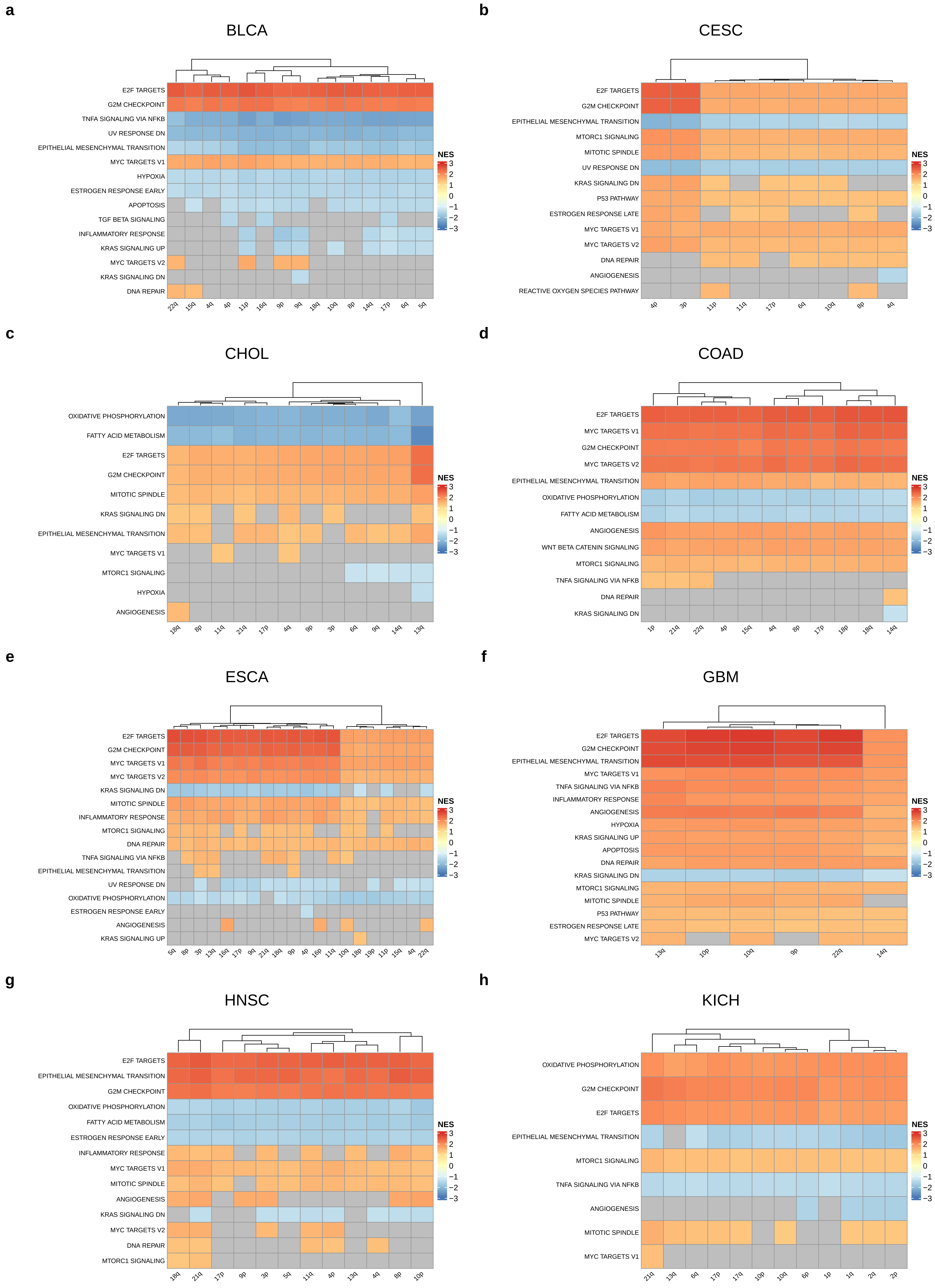


Supplementary Figure 3

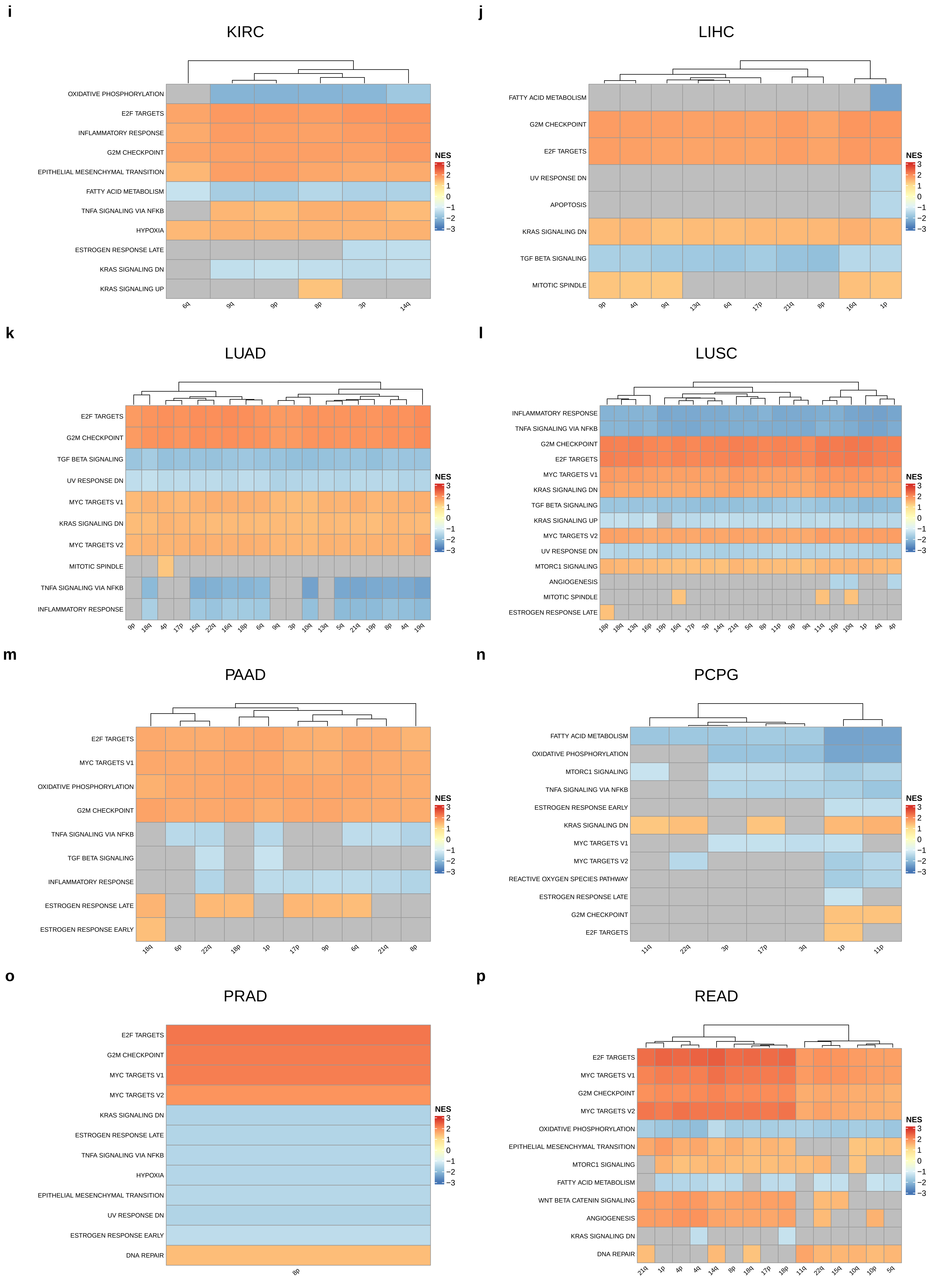


Supplementary Figure 3

**q**

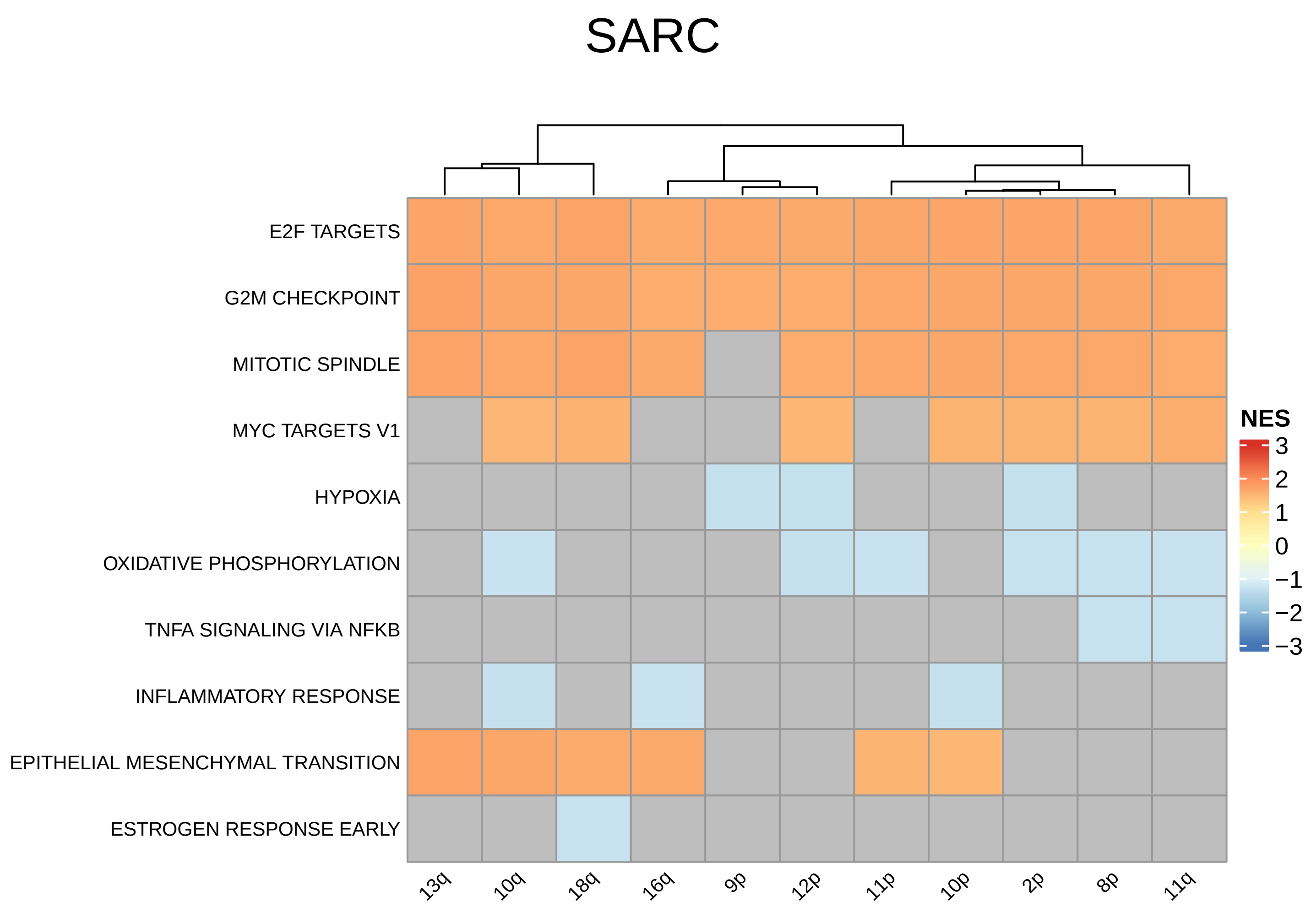


**r**

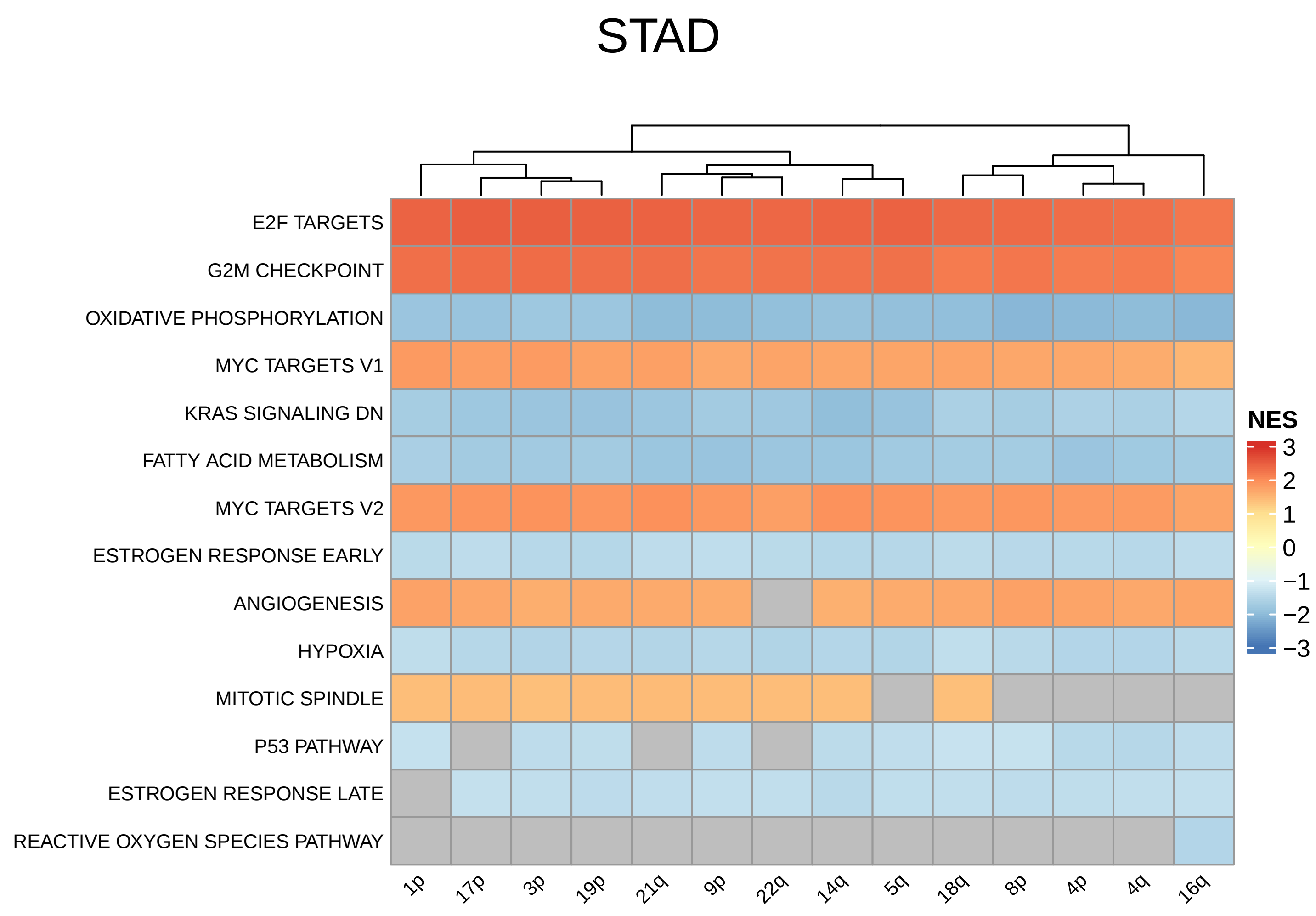


**s**

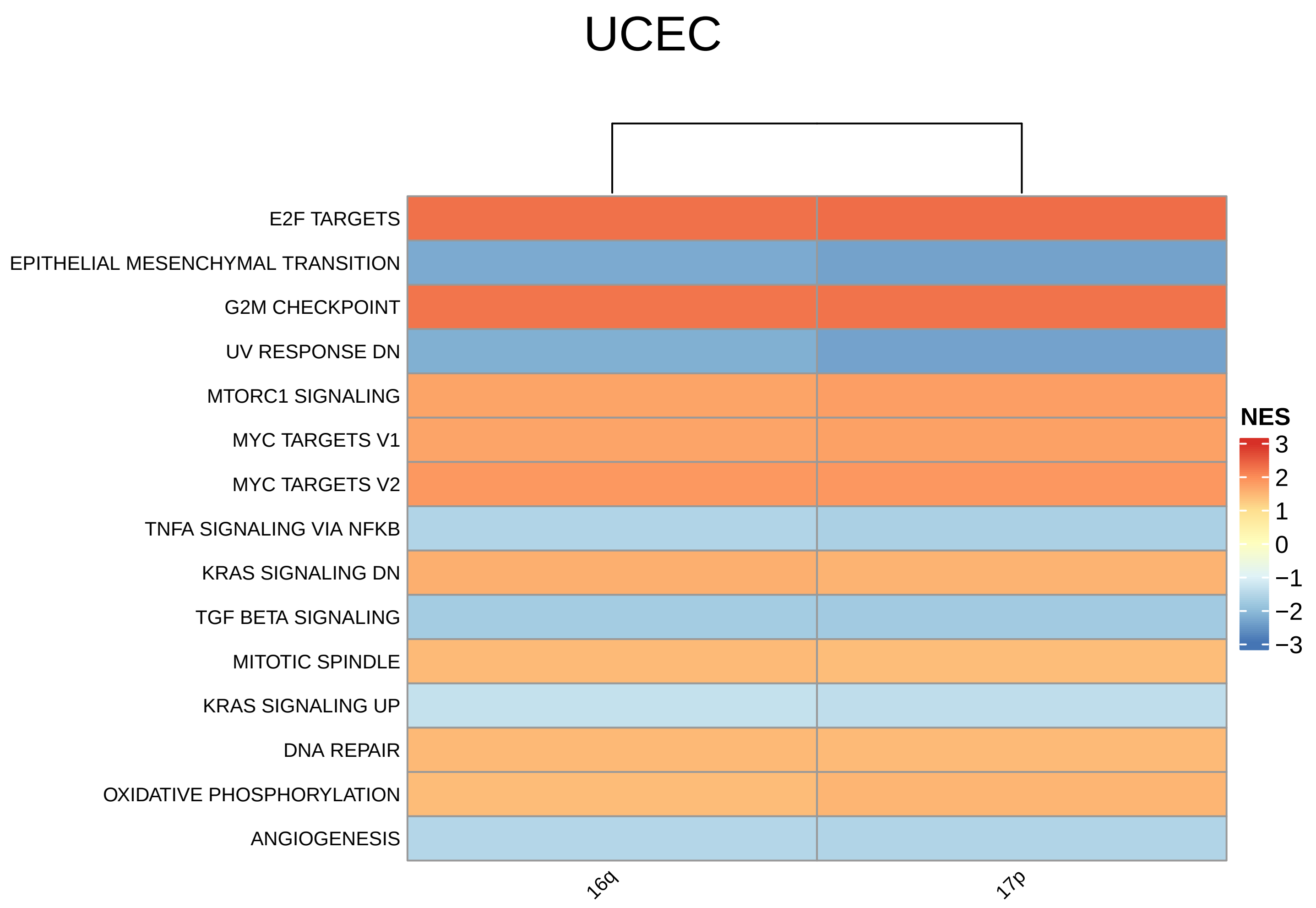


Supplementary Figure 3

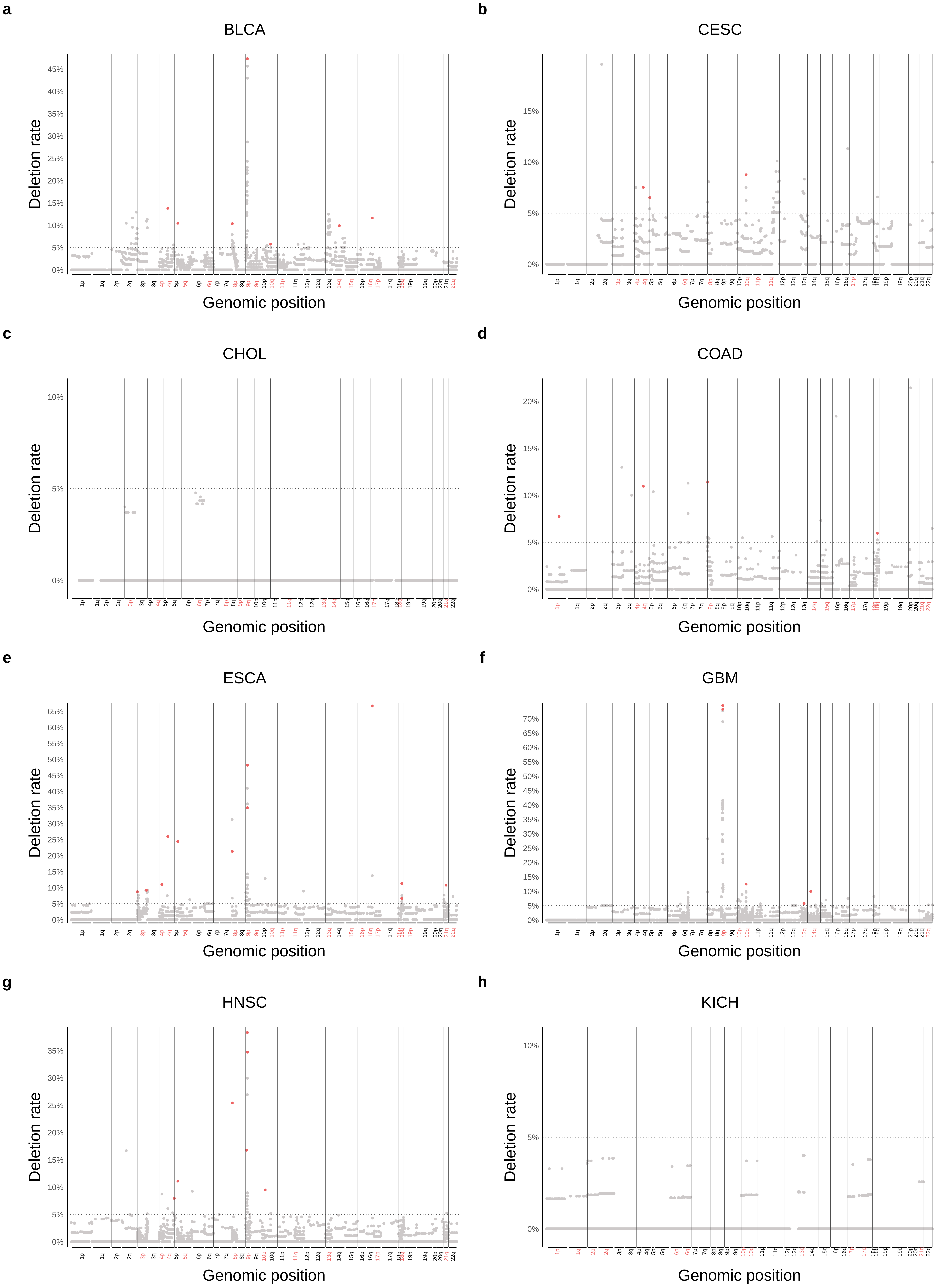


Supplementary Figure 4

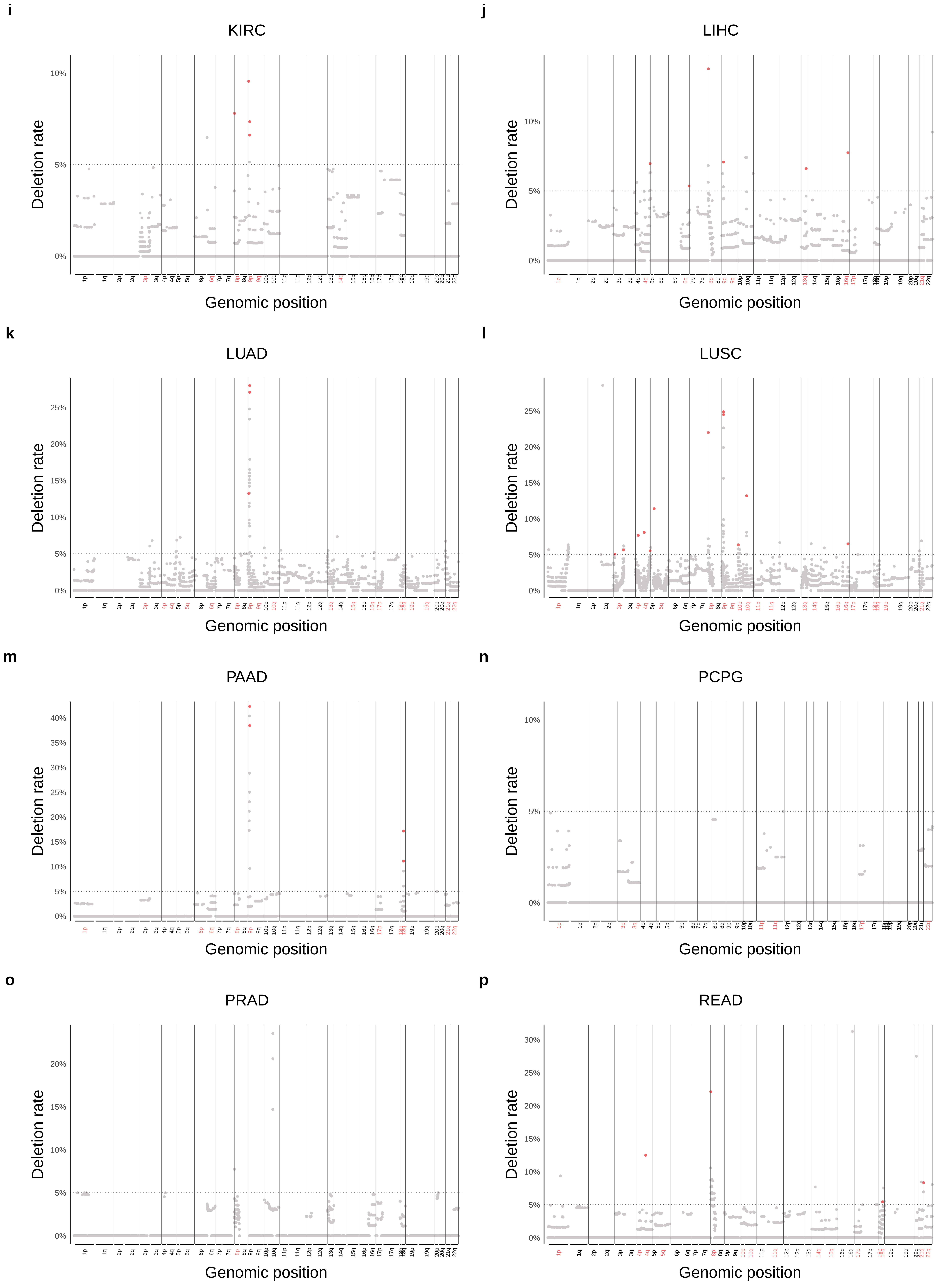


Supplementary Figure 4

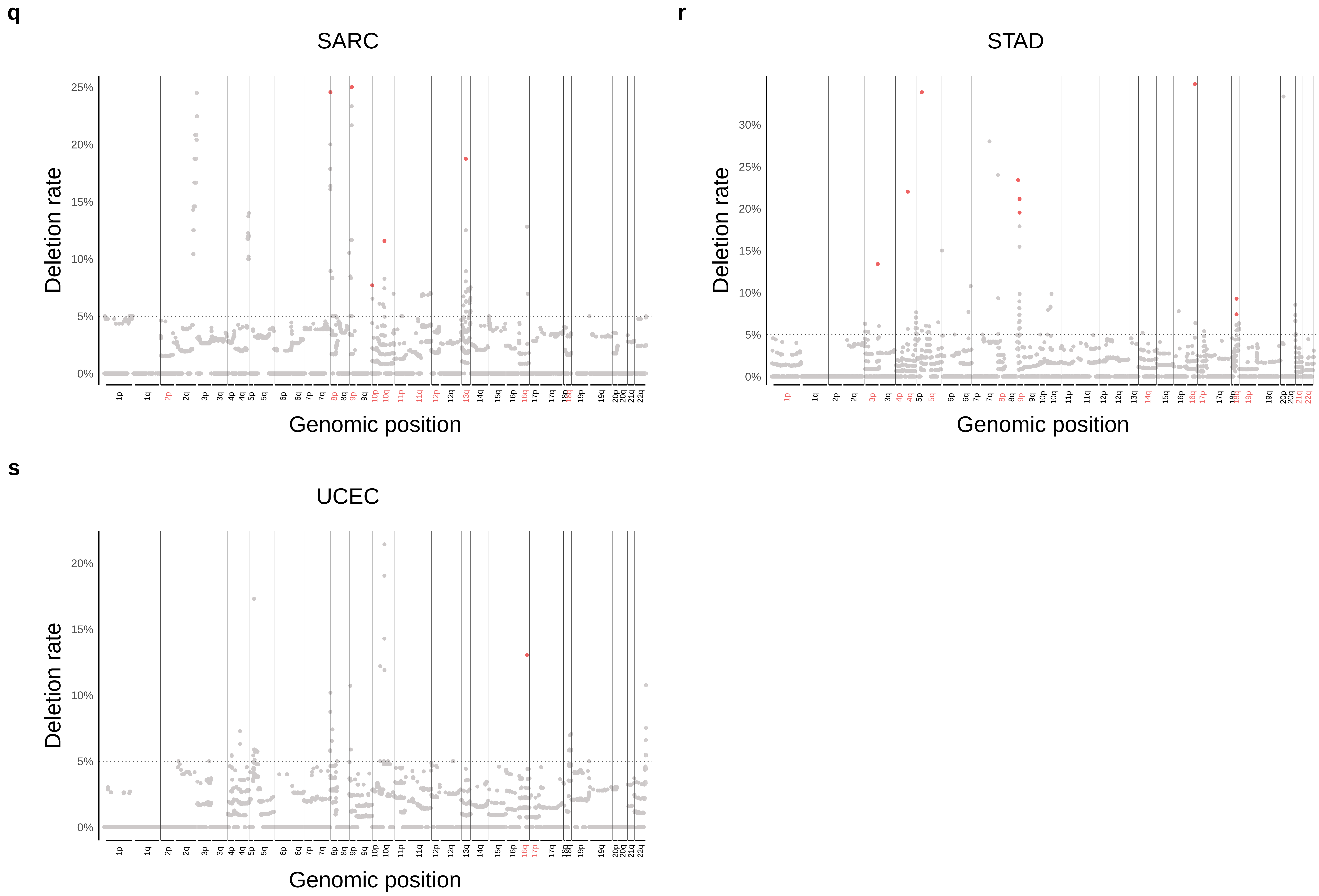


Supplementary Figure 4

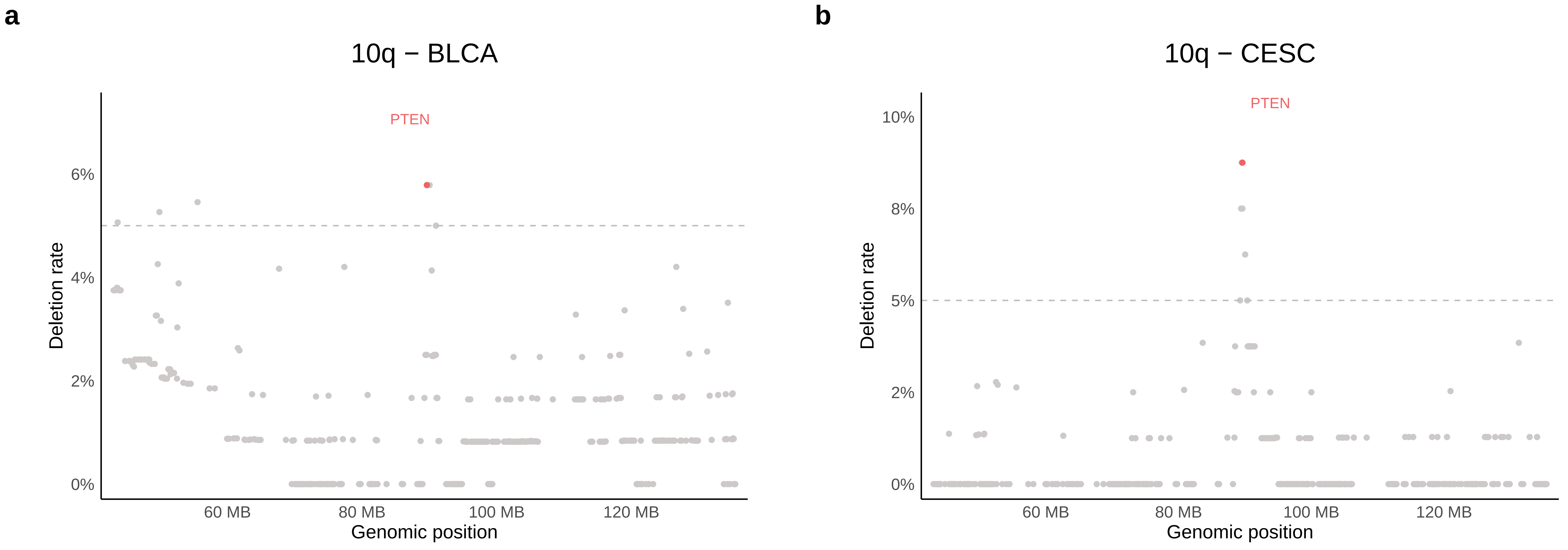


Supplementary Figure 5

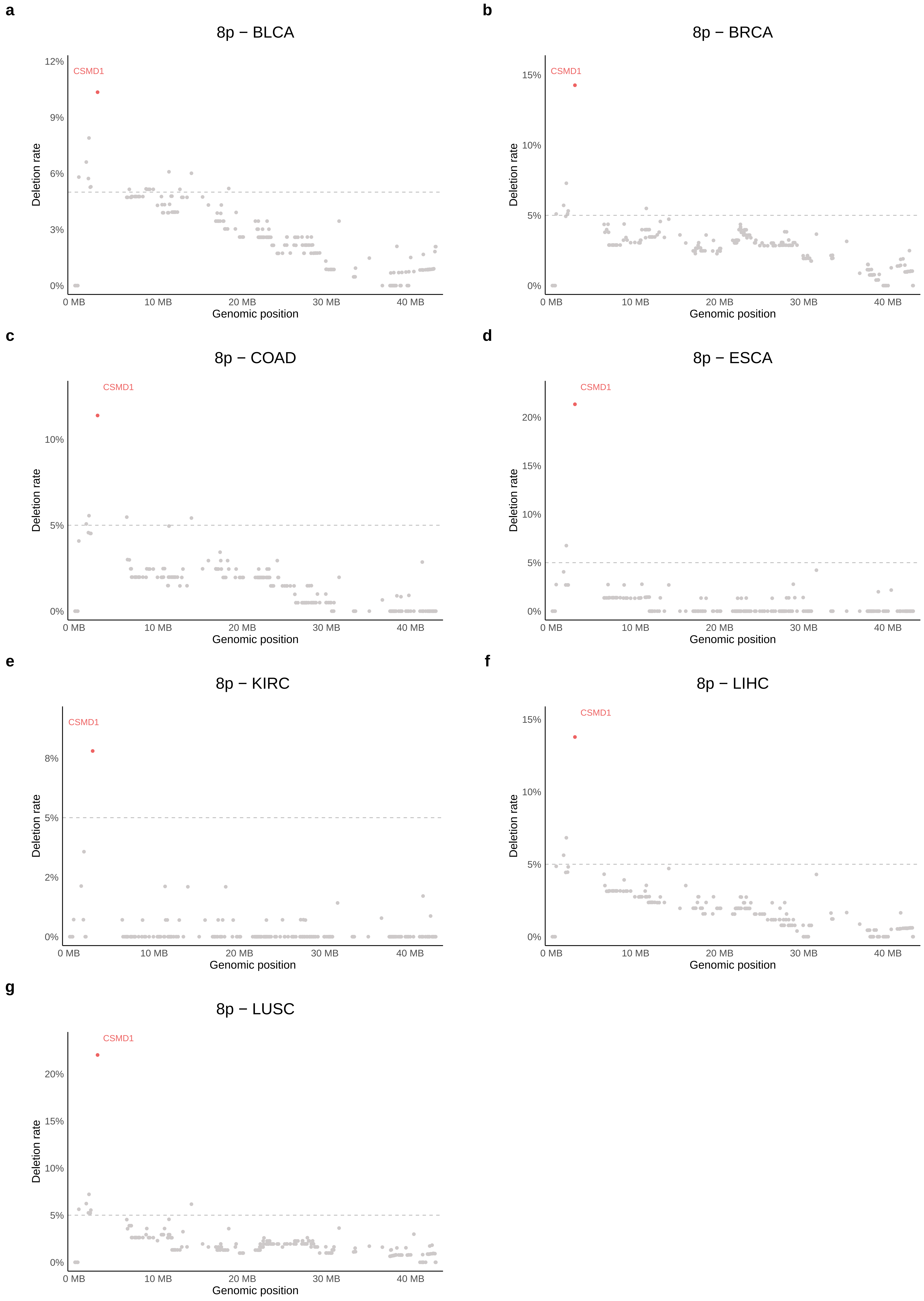


Supplementary Figure 6

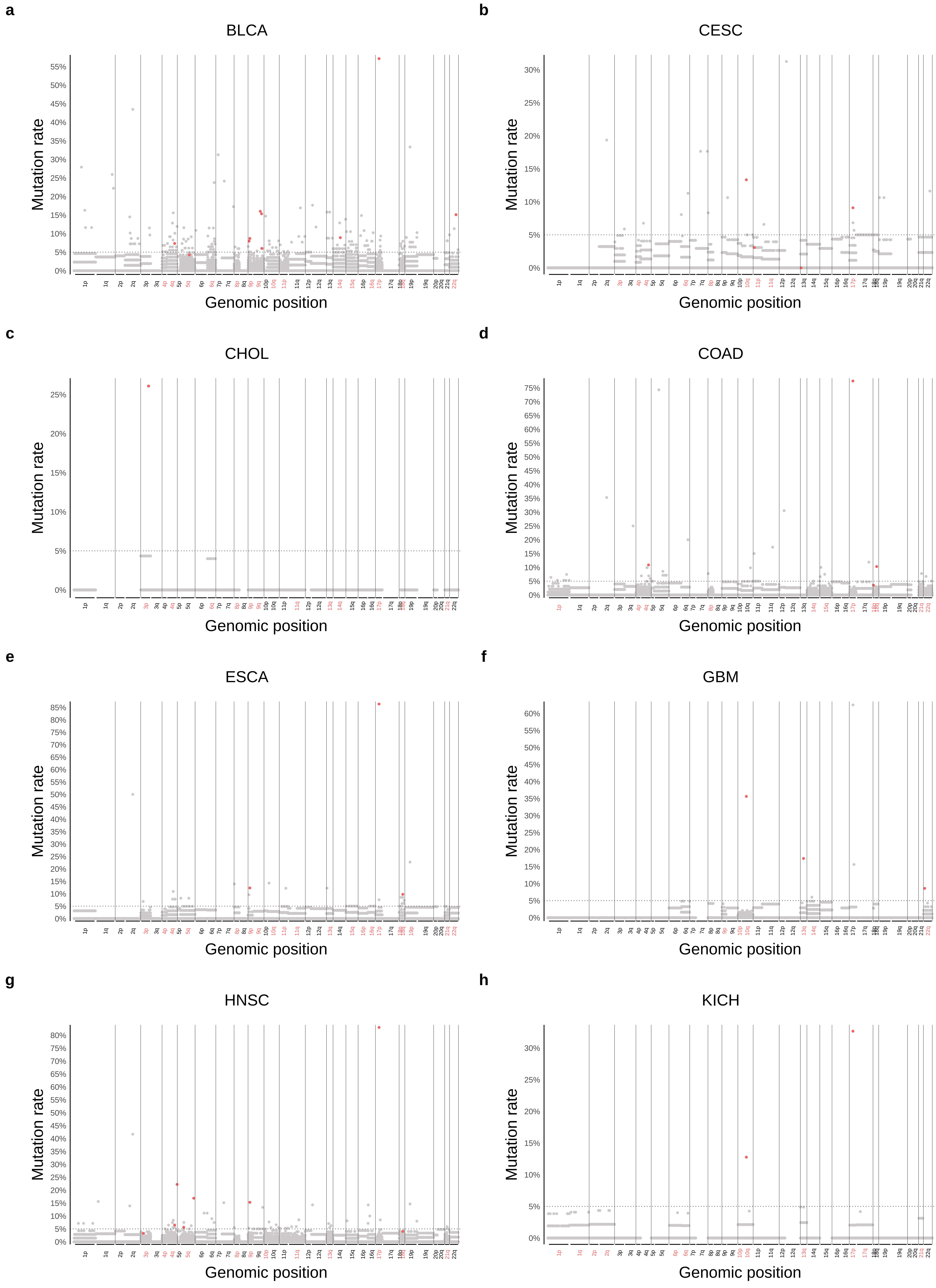


Supplementary Figure 7

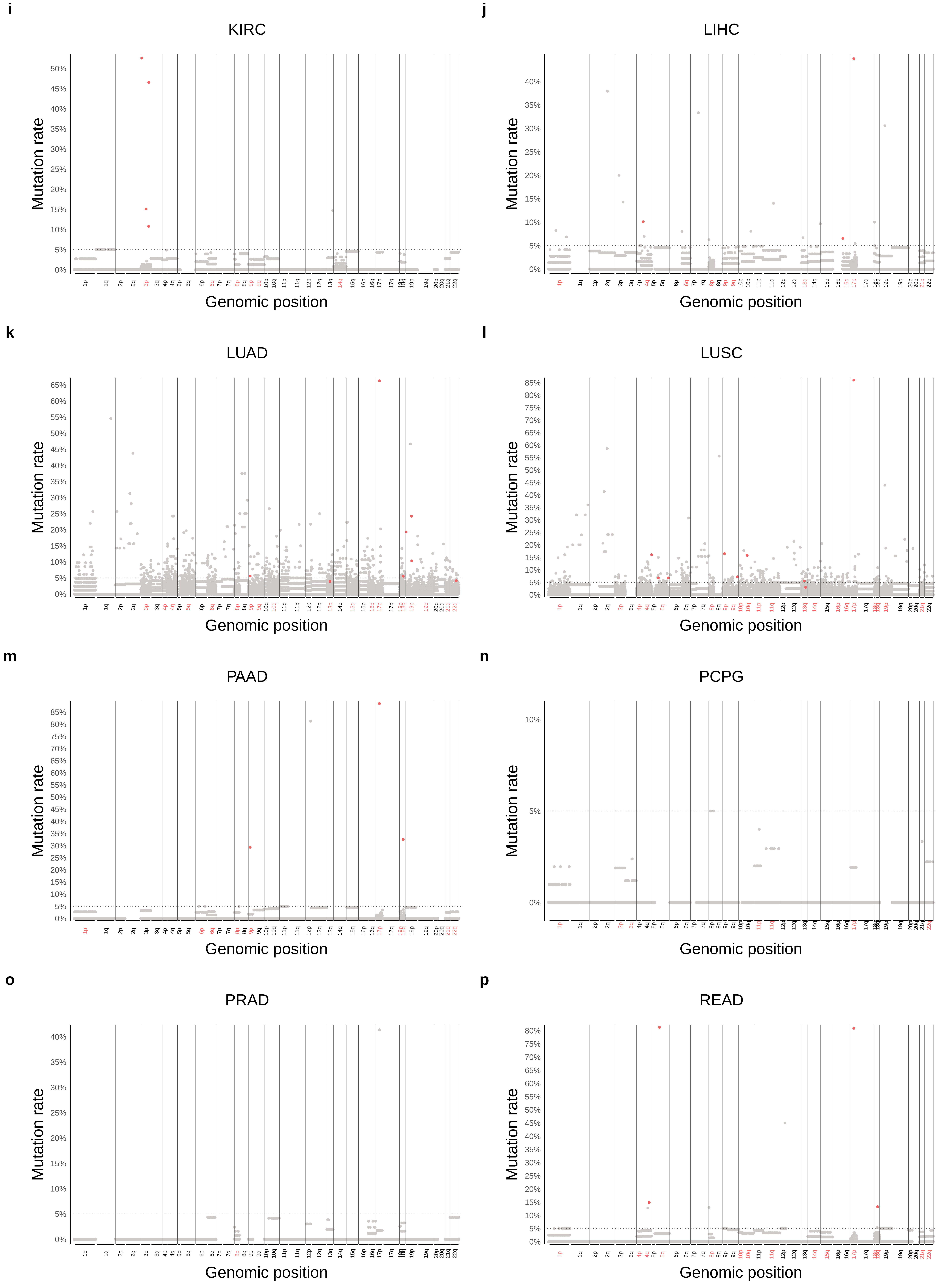


Supplementary Figure 7

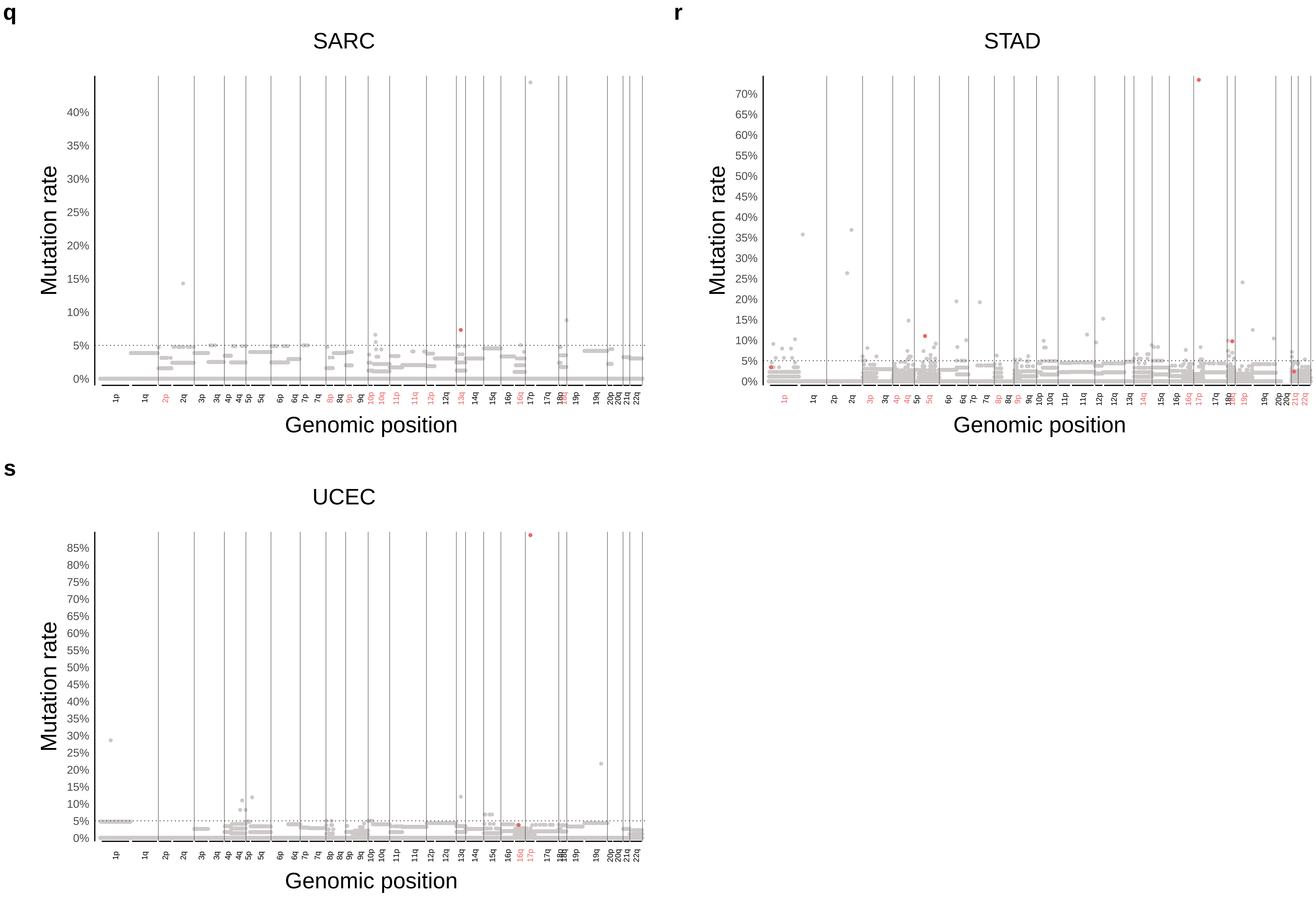


Supplementary Figure 7

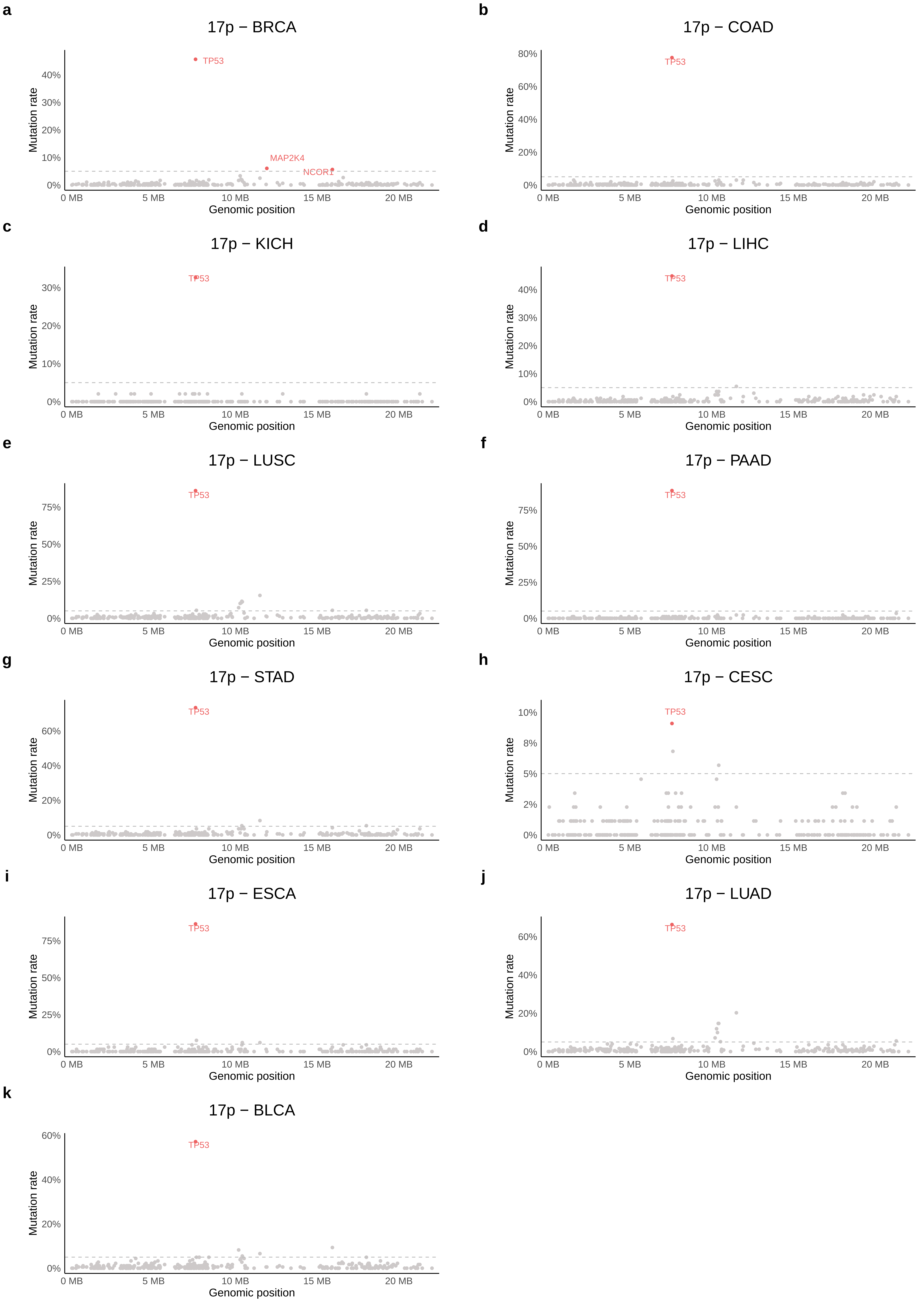


Supplementary Figure 8

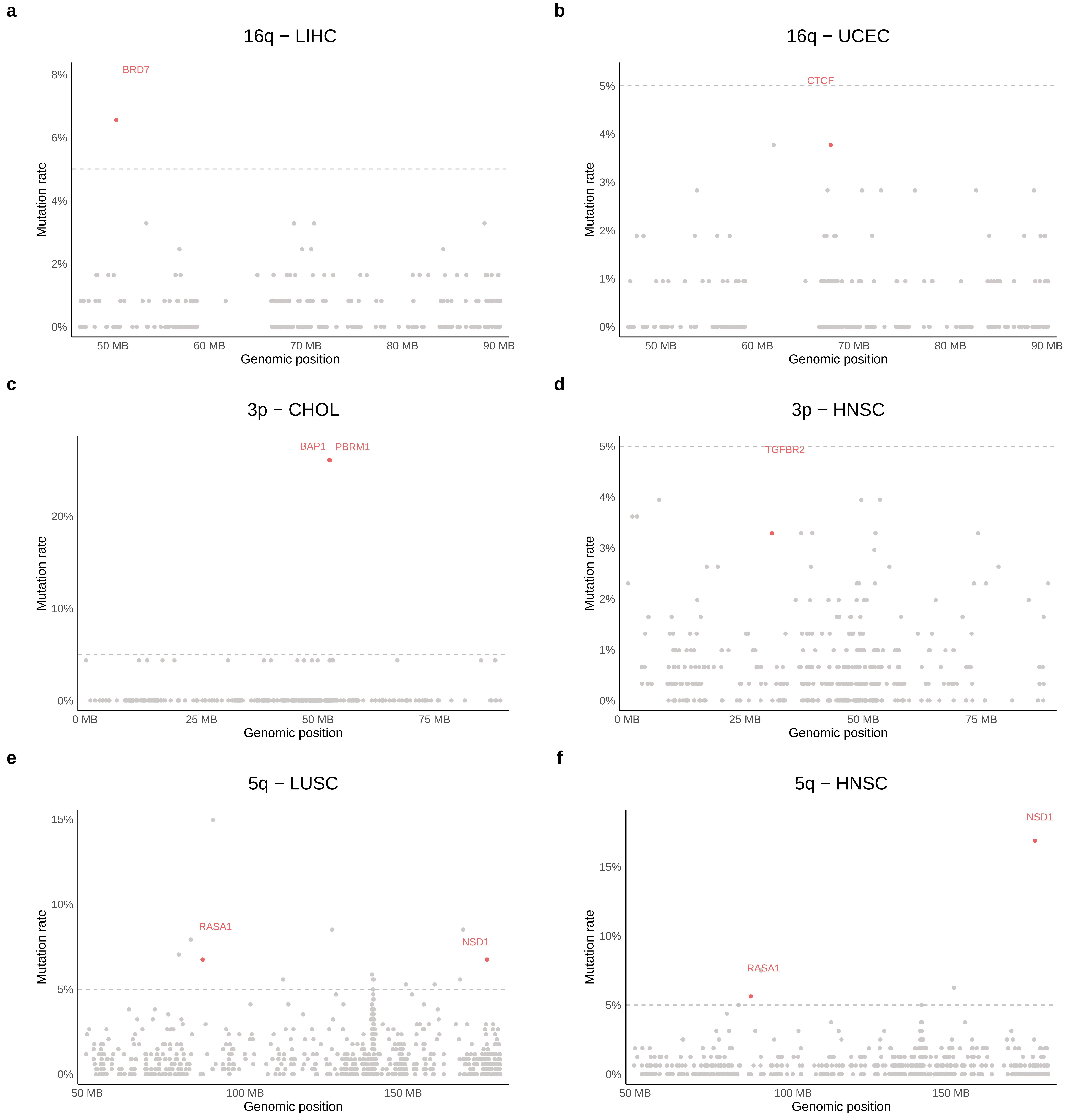


Supplementary Figure 9

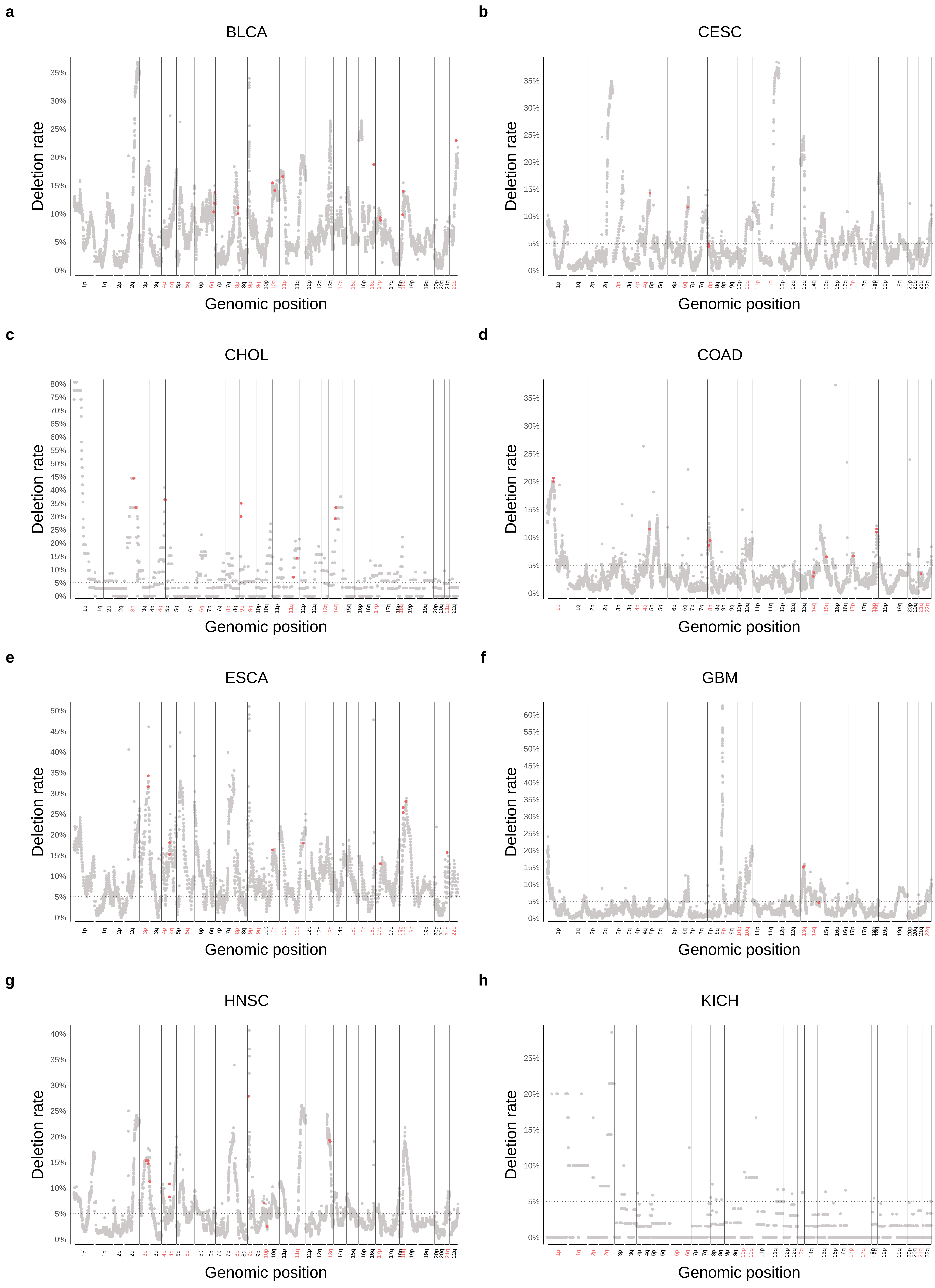


Supplementary Figure 10

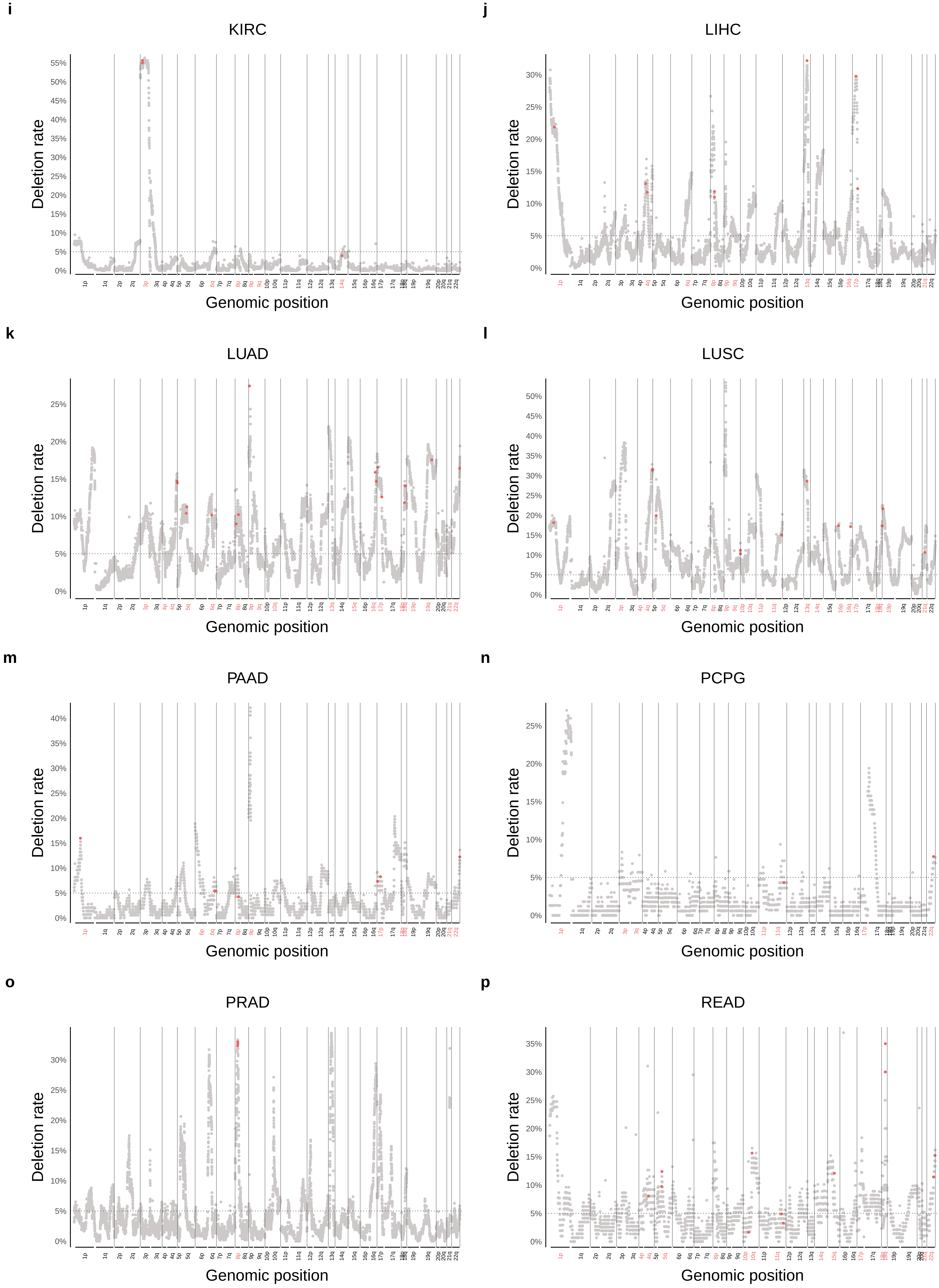


Supplementary Figure 10

q

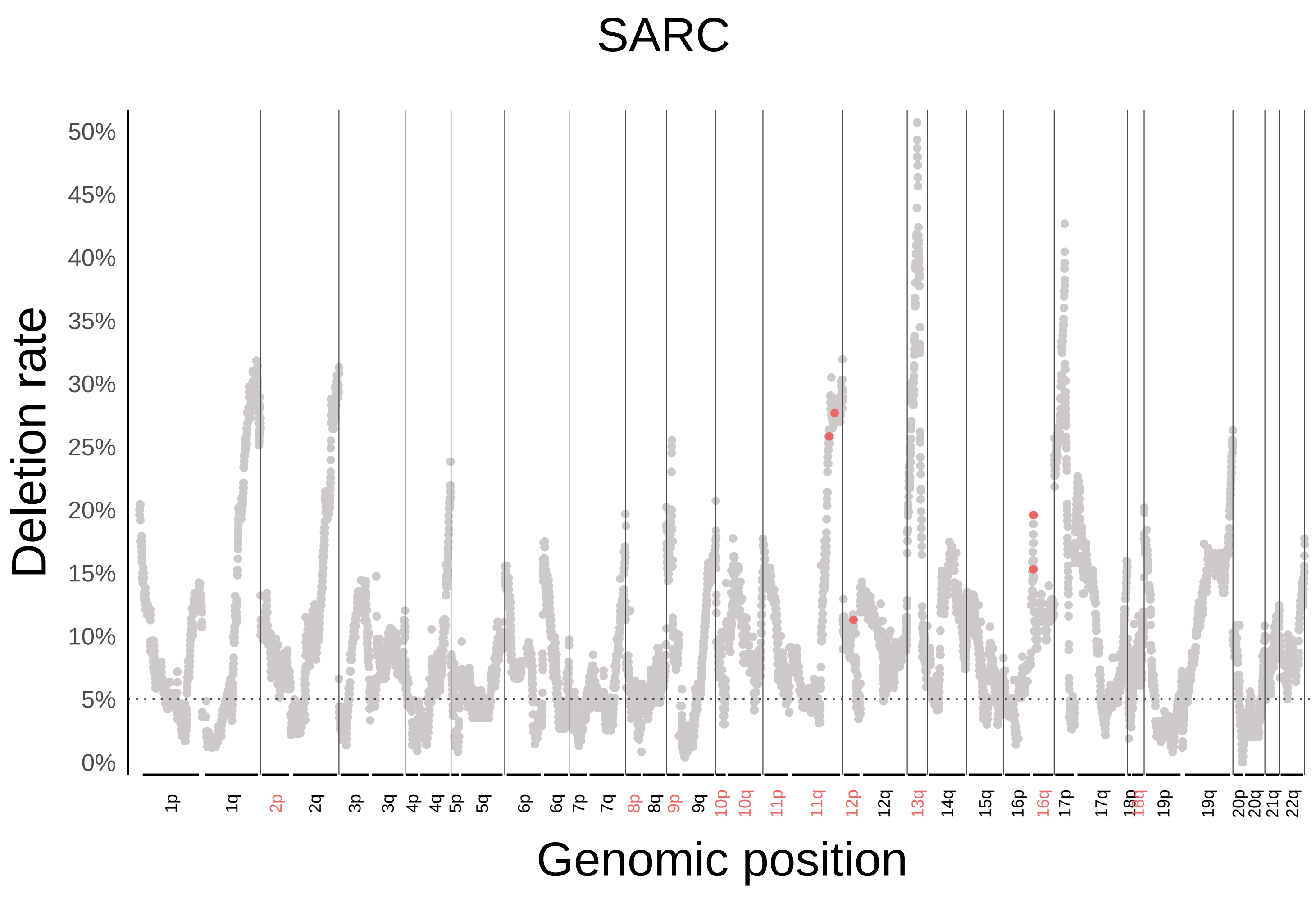


r

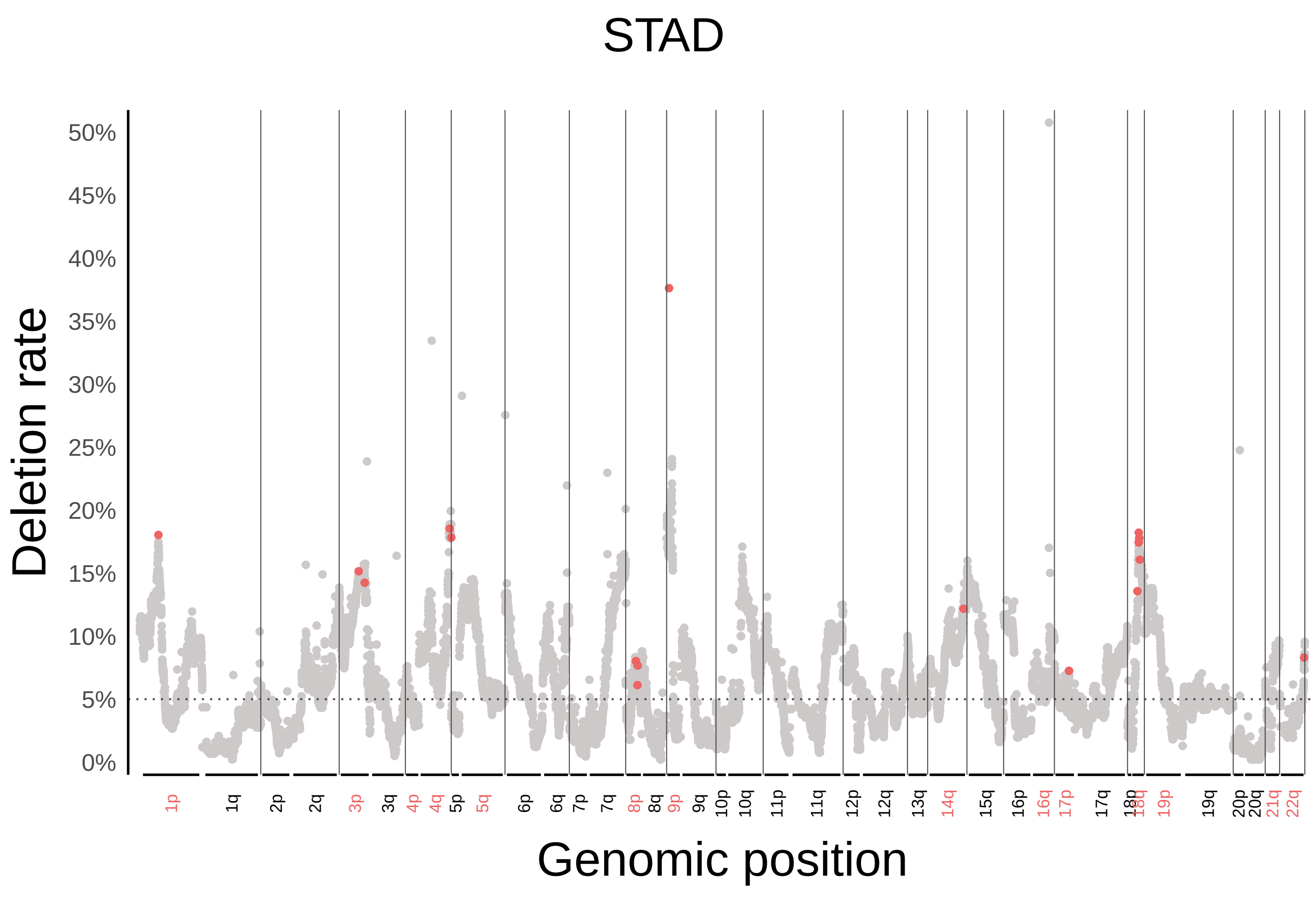


s

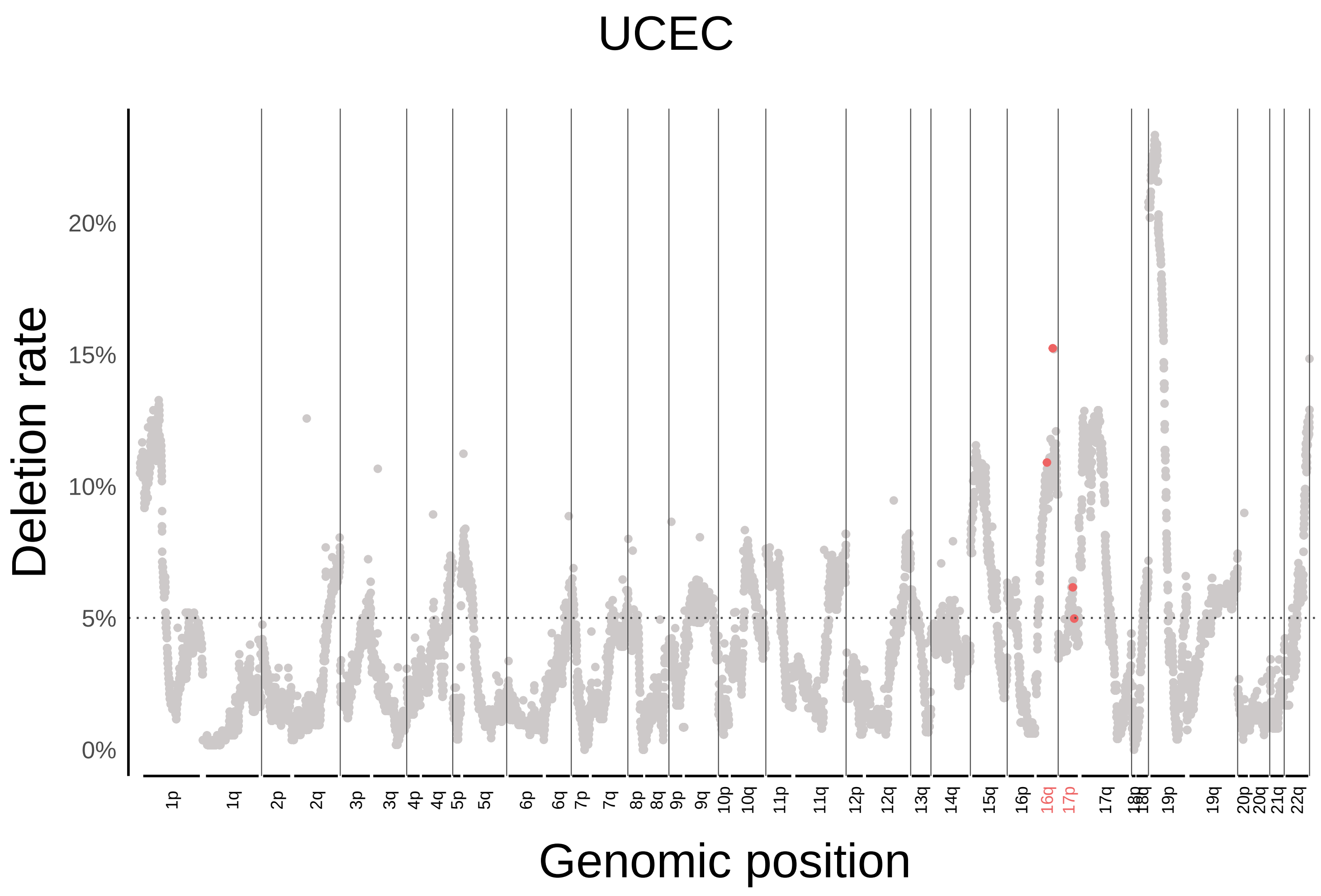


Supplementary Figure 10

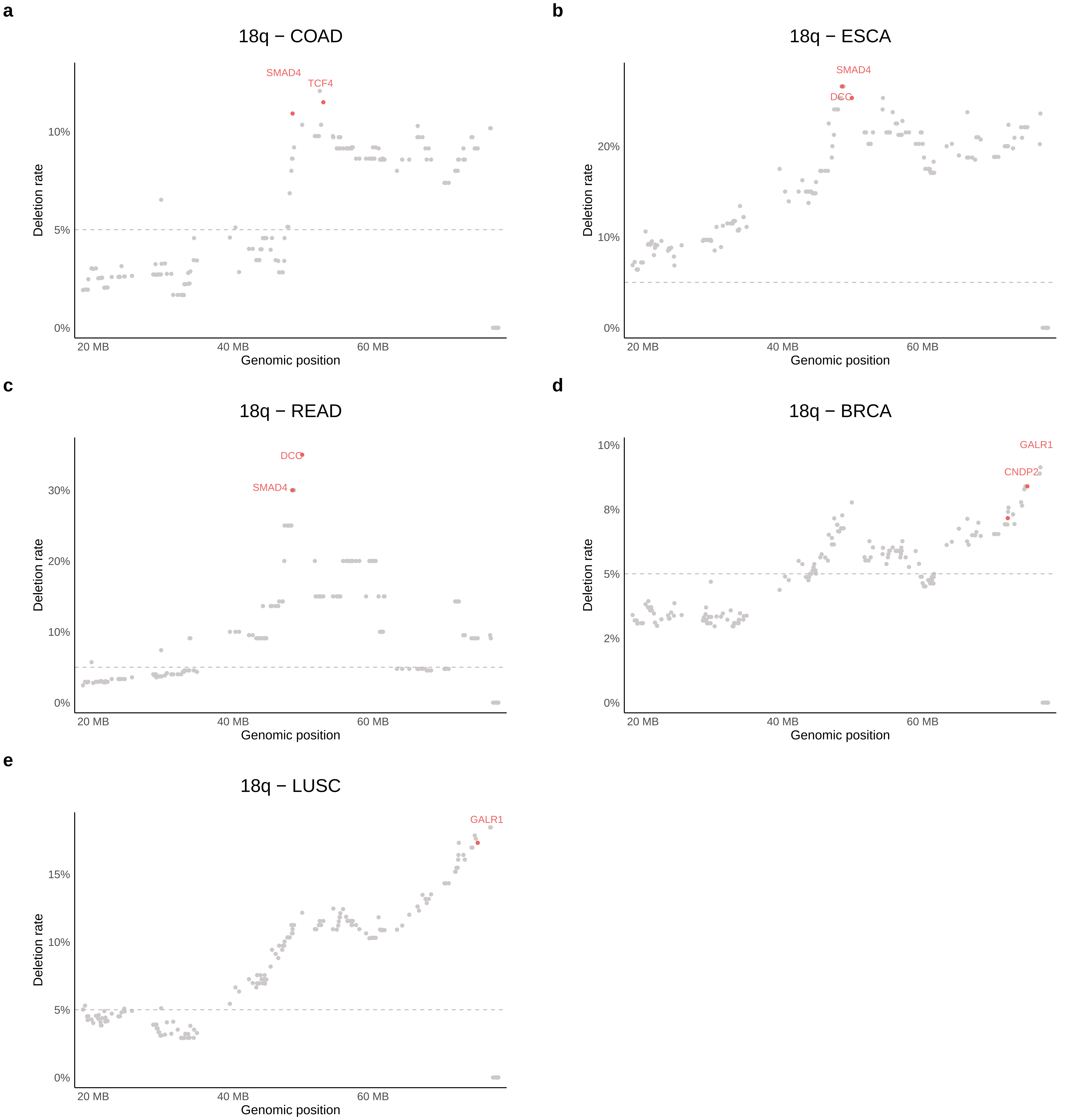


Supplementary Figure 11

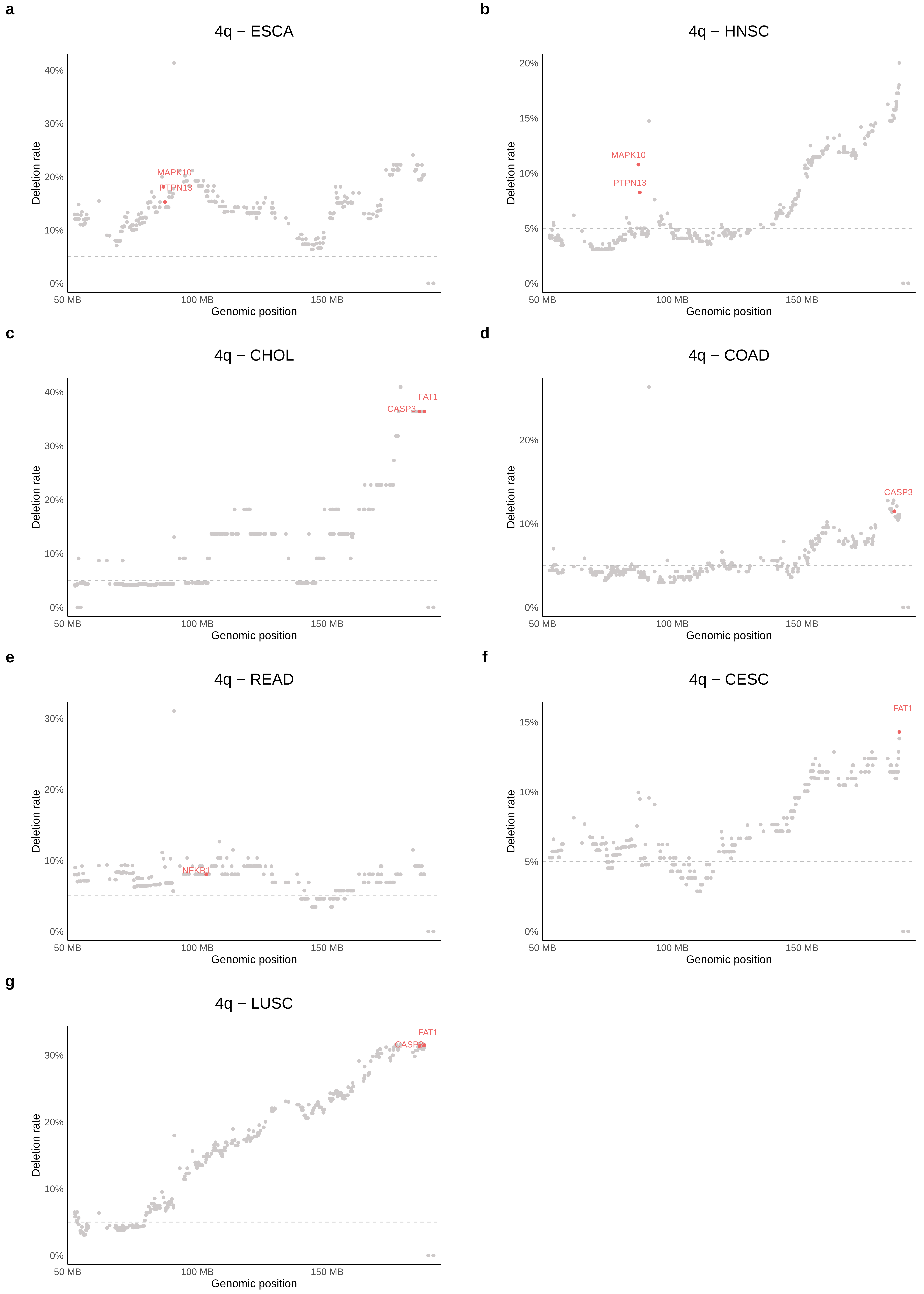


Supplementary Figure 12

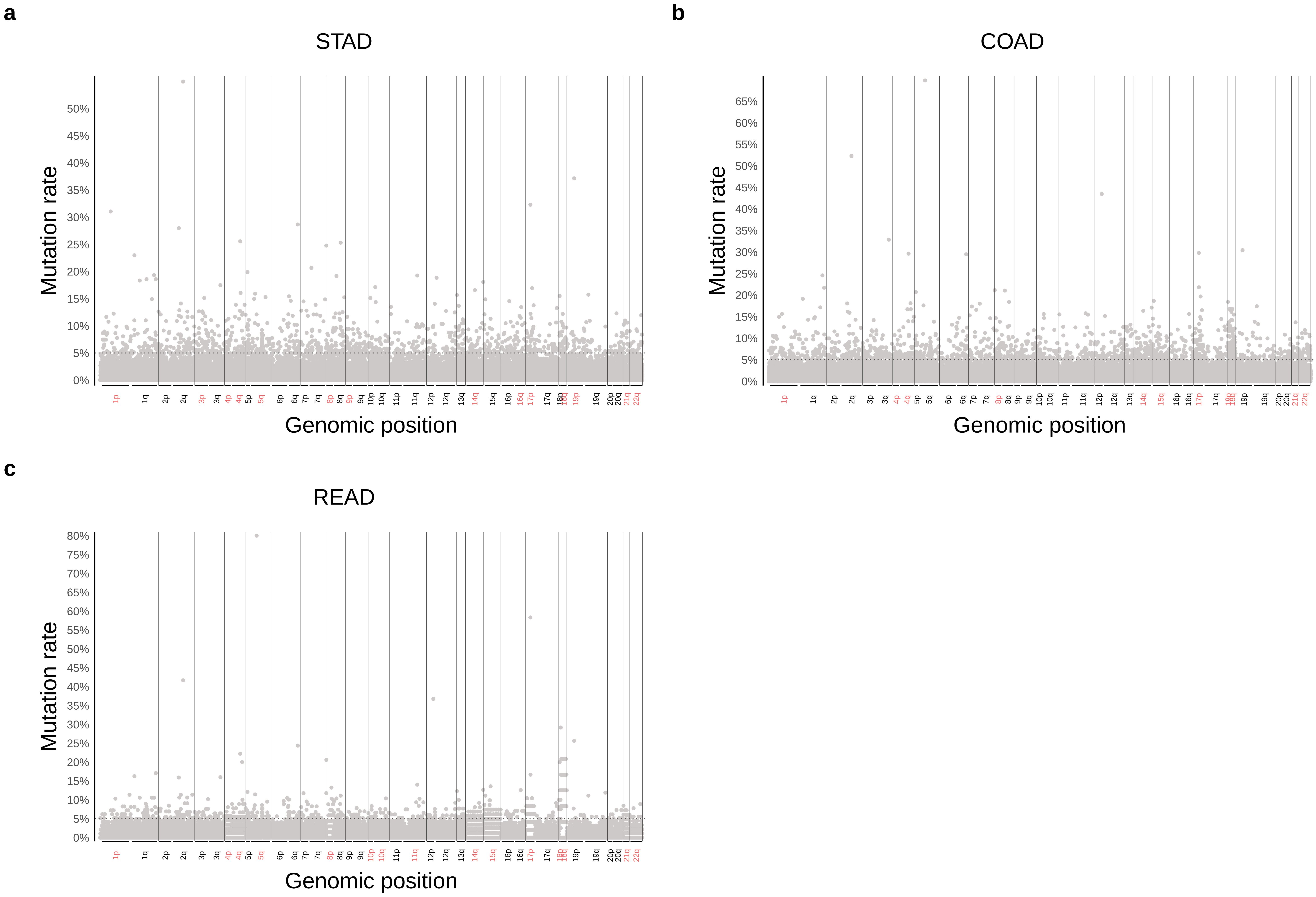


Supplementary Figure 13

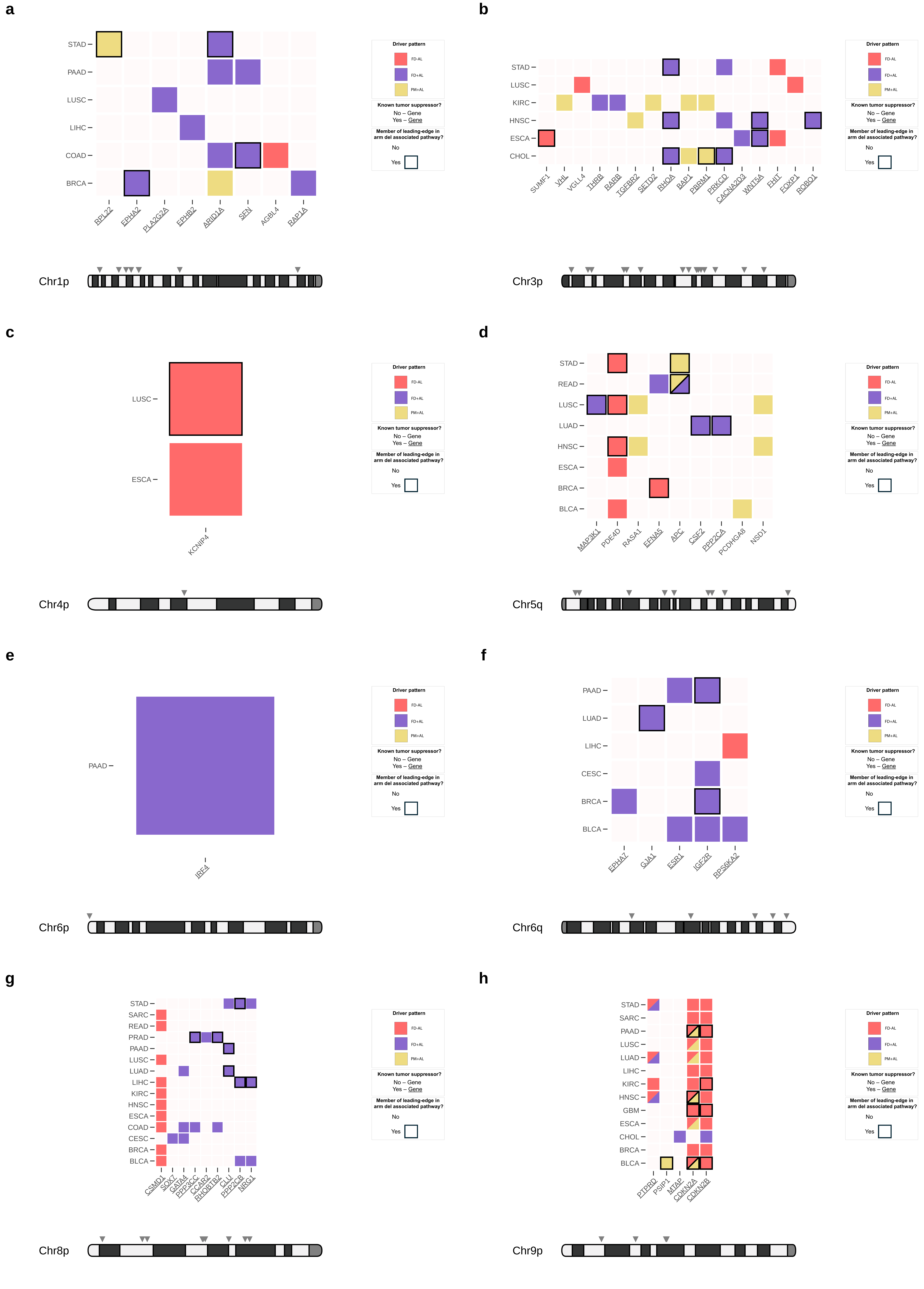

Supplementary Figure 14

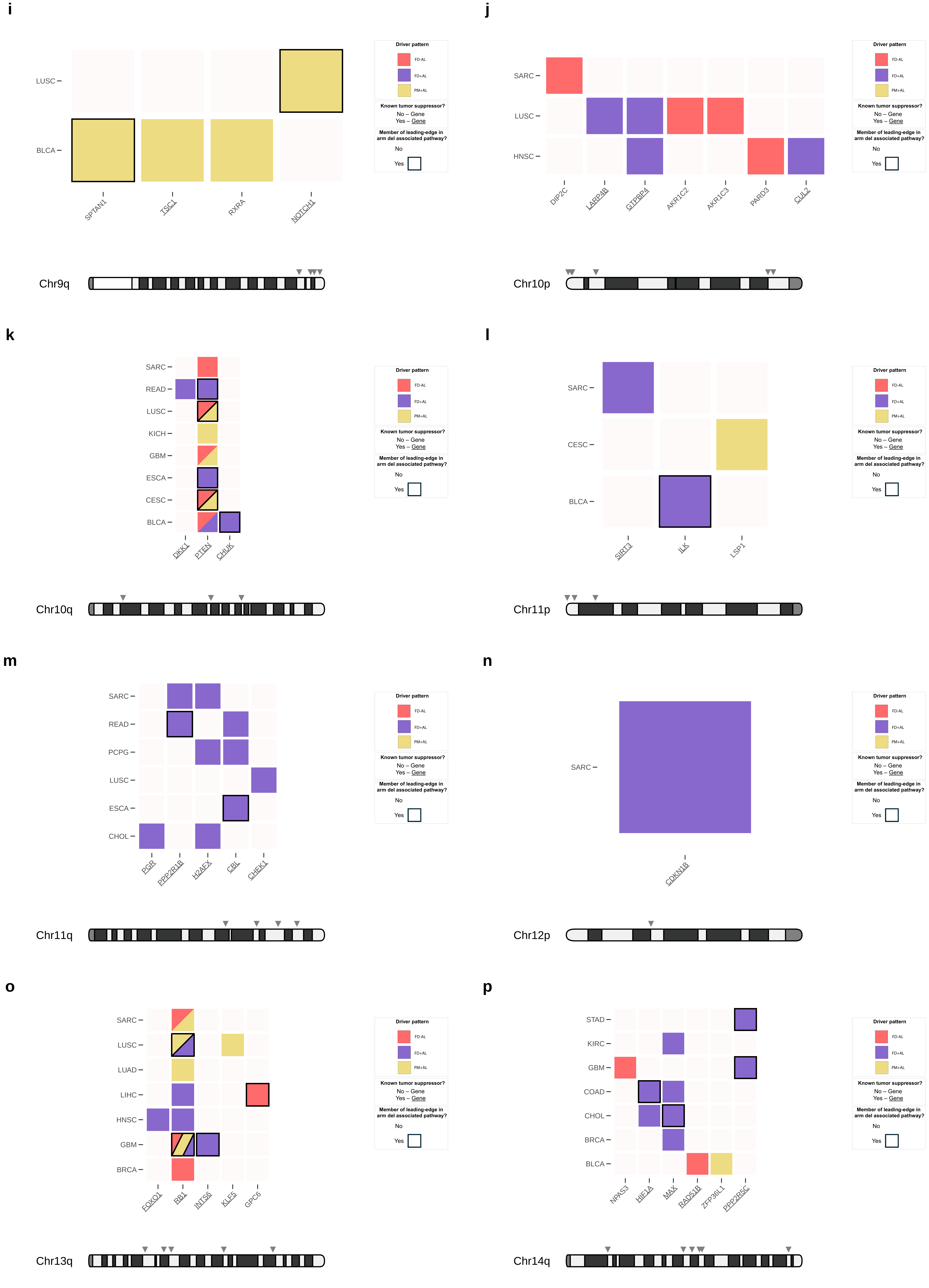


Supplementary Figure 14

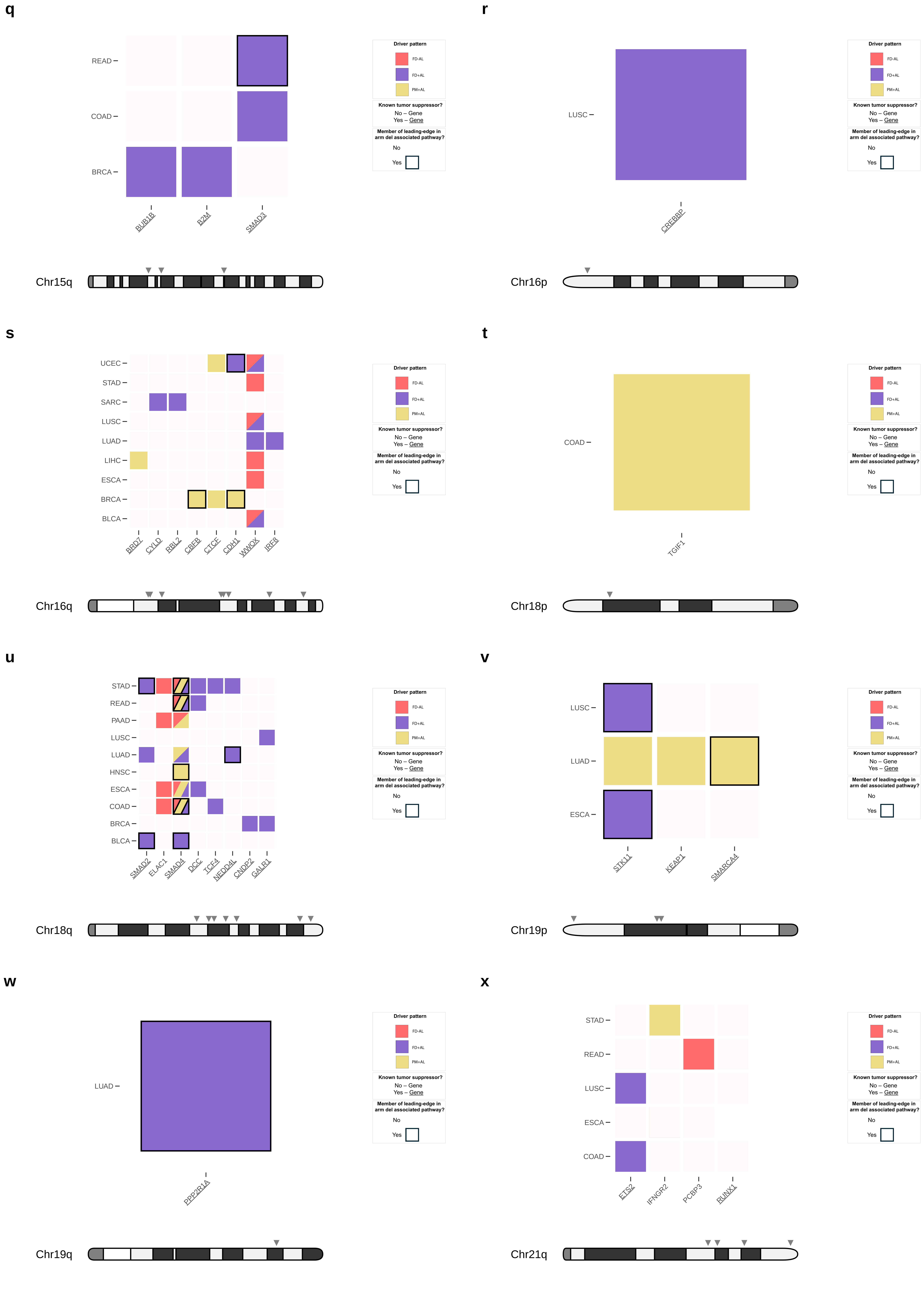


Supplementary Figure 14

y

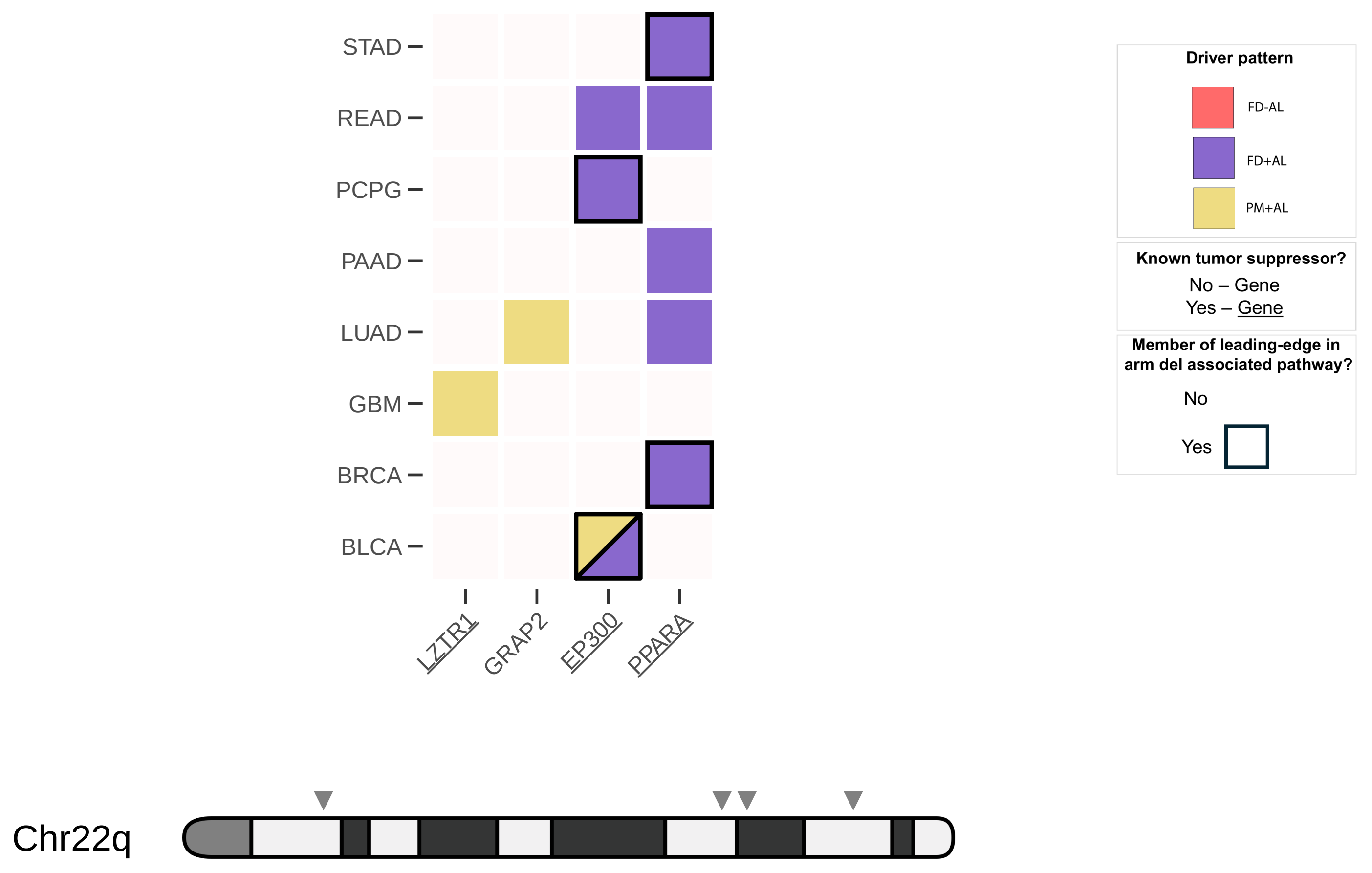
STAD
READ
PCPG
PAAD
LUAD
GBM
BRCA
BLCA
LZTR1
GRAP2
EP300
PPARA
Driver pattern
FD-AL
FD+AL
PM+AL
Known tumor suppressor?
No – Gene
Yes – Gene
Member of leading-edge in arm del associated pathway?
No
Yes
Chr22q
